# Independent signaling pathways provide a fail-safe mechanism to prevent tumorigenesis

**DOI:** 10.1101/2025.02.28.640798

**Authors:** Sari Anschütz, Andrea Schubert, Jobelle M. Peralta, Todd G. Nystul, Katja Rust

**Affiliations:** Institute of Physiology and Pathophysiology, Dept. of Molecular Cell Physiology, Philipps University Marburg, Germany; UCSF, Department of Anatomy, 513 Parnassus Ave, San Francisco, CA 94143, USA; UCSF, Department of OB-GYN/RS, 513 Parnassus Ave, San Francisco, CA 94143, USA; Broad Center of Regeneration Medicine and Stem Cell Research, 513 Parnassus Ave, San Francisco, CA 94143, USA

## Abstract

Controlled signaling activity is vital for normal tissue homeostasis and oncogenic signaling activation facilitates tumorigenesis. Here we use single-cell transcriptomics to investigate the effects of pro-proliferative signaling on epithelial homeostasis using the *Drosophila* follicle cell lineage. Notably, EGFR-Ras overactivation induces cell cycle defects by activating the transcription factors Pointed and E2f1 and impedes differentiation. Hh signaling simultaneously promotes an undifferentiated state and induces differentiation via activation of EMT-associated transcription factors zfh1 and Mef2. As a result, overactivation of Hh signaling generates a transcriptional hybrid state comparable to epithelial-mesenchymal-transition. Co-overactivation of Hh signaling with EGFR-Ras signaling blocks differentiation and induces key characteristics of tumor cells including a loss of tissue architecture caused by reduced expression of cell adhesion molecules, sustained proliferation and an evasion of cell cycle checkpoints. These findings provide new insight into how non-interacting signaling pathways converge at the transcriptional level to prevent malignant cell behavior.

**Graphical abstract:** 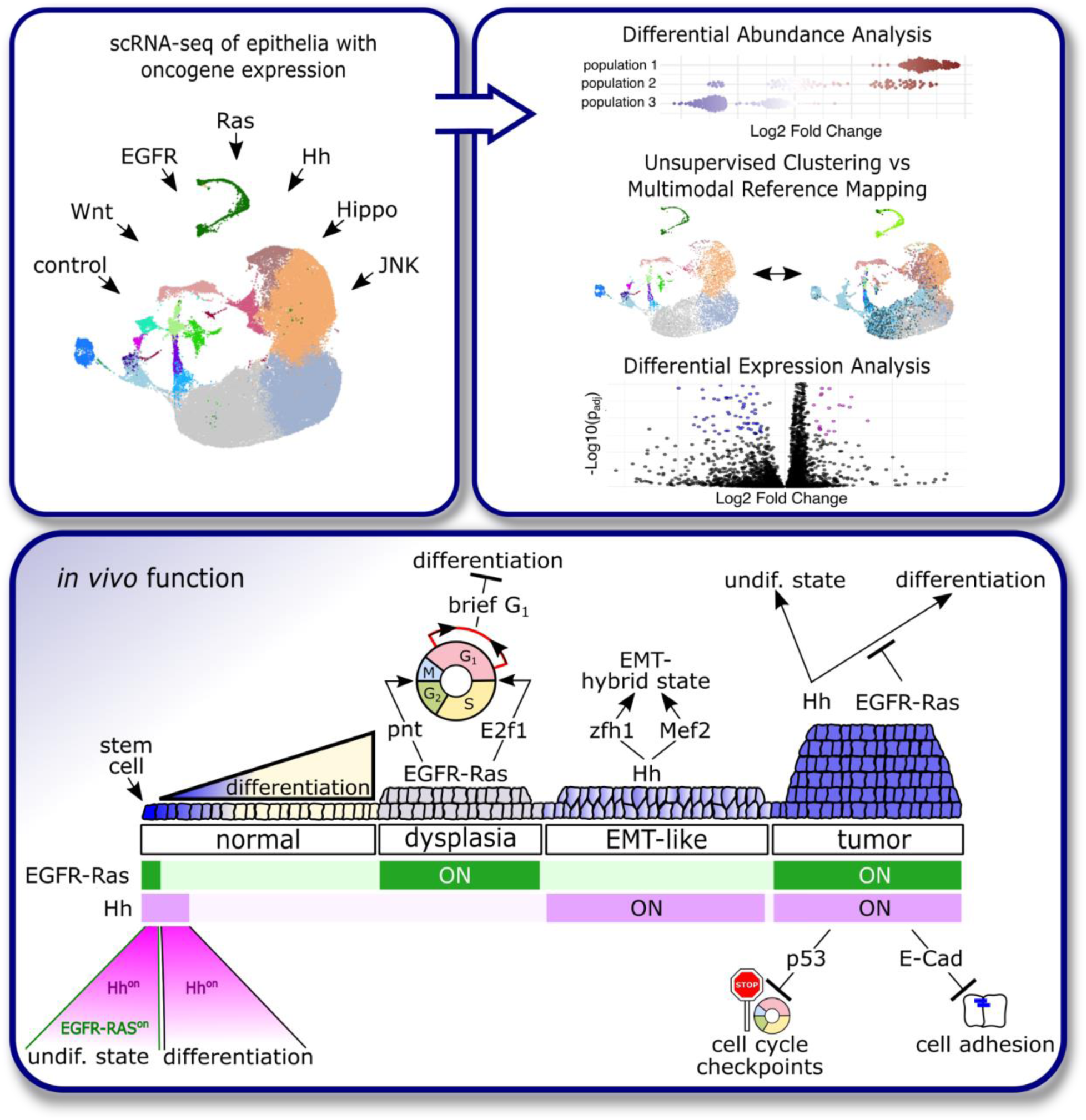

## Introduction

Tissue homeostasis is regulated by the activity of signaling pathways that determine cell behaviors such as the rate of proliferation and the spatial and temporal pattern of differentiation. In epithelial tissues with constitutively active stem cells, these processes occur simultaneously within a relatively small region, thus requiring precise control of each signal. Typically, multiple signaling pathways with unique but overlapping roles function in concert to regulate each cellular behavior. The resulting redundancy and regulatory feedback within this signaling network are thought to confer precision and robustness ^1^. Untangling the contribution of different pathways within the same cells has been a challenge, in part because standard approaches such as immunofluorescence to assay for expression of marker genes or pathway reporters provide information about only a small fraction of the overall transcriptional changes caused by a mutation or other experimental manipulation. In addition, the cellular heterogeneity of most tissues adds additional complexity to the analysis. Signaling pathways that regulate proliferation have also been a major focus of research as they are tightly linked to cancer development ^2^. This includes mutations affecting the activity of the Hedgehog (Hh) and EGFR signaling as well as Ras signaling, which is activated by the EGFR pathway ^3–5^. During cancer development, the Hh and EGFR-Ras pathways can interact and synergistically induce gene expression ^6–8^.

The *Drosophila* follicle stem cell lineage is a well-established model for the study of homeostasis in a stem cell based epithelial tissue ^9^. The Drosophila ovary is composed of long strands of developing follicles, called ovarioles, and the follicle epithelium is a layer of somatic cells that surrounds each follicle. The follicle epithelium is maintained by a population of follicle stem cells (FSCs) that reside at the anterior edge of the tissue in a structure called the germarium. The precise number and location of the FSCs has been debated recently ^10,11^, but it is generally agreed that they are located between Regions 2a and 2b of the germarium at or near the anterior boundary of the expression domain of the cell surface marker, Fasciclin 3 (Fas3). FSCs produce transiently amplifying daughter cells, called prefollicle cells (pFCs) (Fig. 1A) that differentiate into either main body (MB) follicle cells, which ensheath the growing germline cysts, stalk cells, which connect adjacent cysts, or polar cells on either end of the cysts which regulate MB patterning and stalk cell differentiation (Fig. 1A). Previous studies using genetics, lineage tracing, and single-cell transcriptomics have revealed that differentiation toward the polar, stalk, or MB follicle cell fate is initiated early in the FSC lineage, at a stage that is well before any morphological distinctions in the pFC population are apparent ^12–15^. At this stage, multiple signaling pathways function together to ensure that each pFC differentiates into the correct cell type during the right time and place. The EGFR-Ras and Hedgehog (Hh) signaling pathways are all required for FSC self-renewal and are important regulators of follicle cell proliferation and differentiation ^9,16–18^. EGFR signaling, which is indirectly activated by Wnt signaling in FSCs, functions upstream of both the Ras/MAPK and the Lkb1/AMPK pathways to promote FSC self-renewal, regulate FSC and pFC proliferation, and establish apical polarity in the early follicle cell lineage ^16^. Hh signaling independently controls proliferation and differentiation in the FSCs and early pFCs (Fig. 1A). Hh signaling induces proliferation via the activation of Hippo signaling and regulates differentiation in part through direct and indirect inhibition of the transcription factors, castor (cas) and eyes absent (eya) ^14,19–22^.

**Figure 1:**
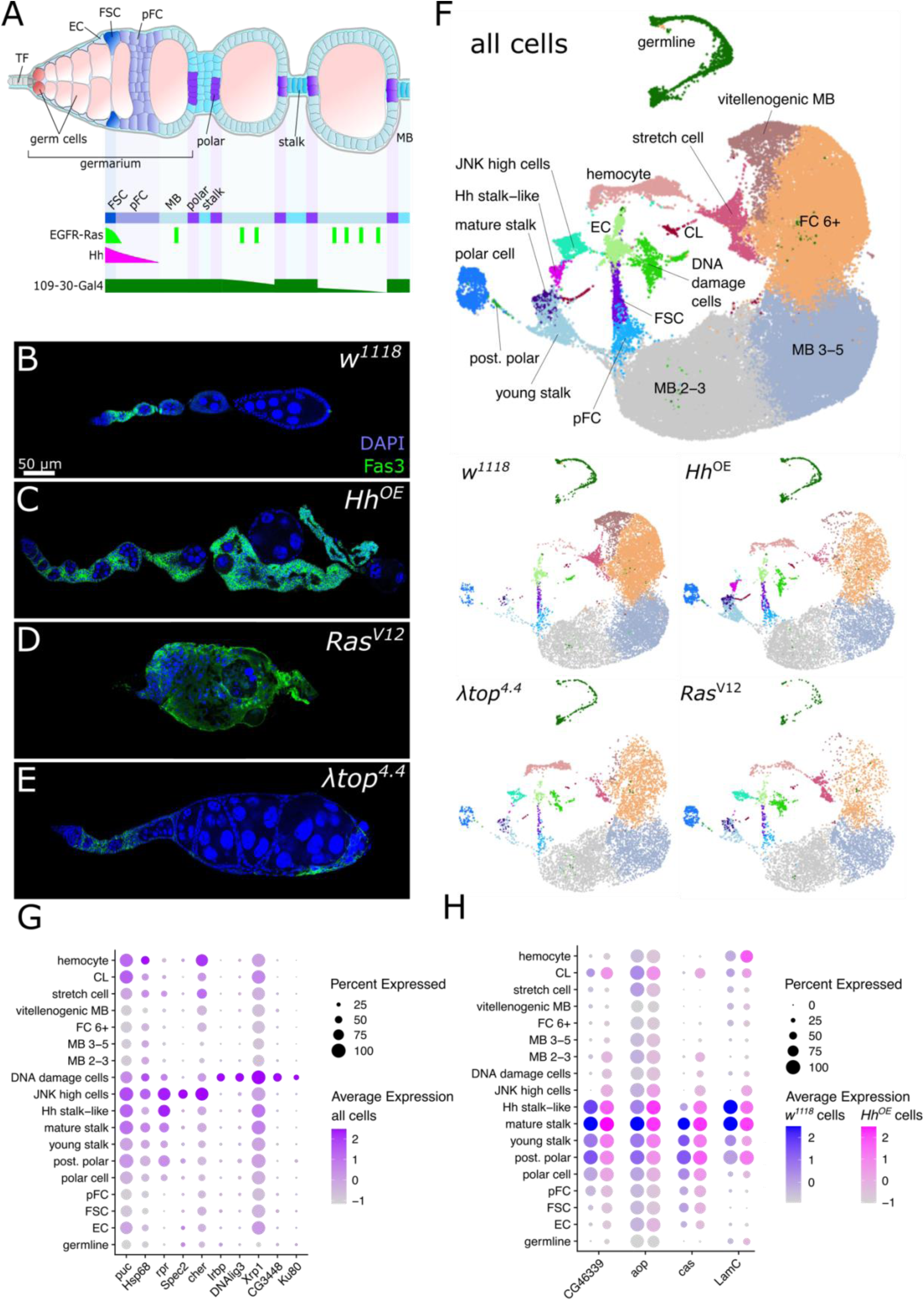
Single-cell RNA-seq of follicle cells with overactivated signaling pathways. A) Graphic depiction of a *Drosophila* germarium and early stages of oogenesis and activity of EGFR-Ras, Hh signaling and the 109-30-Gal4 driver. TF: terminal filament, EC: escort cell, FSC: follicle stem cell, pFC: prefollicle cell, MB: main body follicle cell. B-E) Ovarioles of 109-30^ts^ crossed to the respective alleles and stained for DAPI (blue) and Fas3 (green). F) Cluster identity represented on a UMAP plot of ovarian cells from all genotypes (all cells) or separate genotypes induced by 109-30^ts^ as indicated. G) Percent and average expression of markers of the clusters “JNK high cells” and “DNA damage cells” in all cells. H) Percent and average expression of stalk cell markers in *w^1118^* and *Hh^OE^* cells.

Here, we apply single-cell RNA-sequencing, bioinformatic analyses, and *in vivo* assays to investigate ectopic activation of the pro-proliferative pathways, Wnt, EGFR-Ras, JNK, Hh and Hippo, signaling using the *Drosophila* follicle epithelium as a model. We find that co-activation of Hh and EGFR-Ras signaling pathways induces follicle cell tumors and reveal the transcriptional landscapes of distinct steps in tumor development from the healthy epithelium to a precursor state in which either Hh or EGFR-Ras are overactivated, to the cancer stage where both pathways are overactivated. While overactivation of EGFR-Ras results in increased proliferation and cell cycle anomalies, Hh overactivation induces an undifferentiated state while concurrently promoting differentiation towards the stalk cell fate. This induces a transcriptional hybrid state in cells predetermined to differentiate towards polar or MB cell fates, while undifferentiated FSCs and pFCs are protected from differentiation. Co-overactivation of Hh and EGFR-Ras signaling pathways results in a massive expansion of the pool of transiently amplifying pFCs. This suggests that Hh activation induces differentiation only in the absence of active EGFR-Ras signaling. These hypertrophic pFCs undergo multilayering by misregulation of cell adhesion molecules and evade cell cycle checkpoints, thereby displaying major hallmarks of cancer.

## Results

### Mapping pro-proliferative signaling pathways at single-cell resolution

To identify the transcriptional targets of the pro-proliferative signaling pathways in the follicle cell lineage, we used the early follicle cell driver, 109-30-Gal4 ^23^ and a temperature-sensitive Gal80 (*Gal80^ts^*) ^24,25^ to constitutively activate pro-proliferative pathways specifically during adulthood and then performed single-cell RNA-sequencing (Fig. 1A). We selected UAS-driven alleles that produced strong morphological phenotypes that are consistent with previous studies (Fig. 1B-E, S1A-C) ^16,17,26–28^. Specifically, we expressed the wildtype Hh ligand to overactivate Hh signaling (Hh^OE^) ^29^ and a constitutively active allele of EGFR (EGFR^λtop4.4^, hereafter referred to as λtop^4.4^) ^30^ to overactivate EGFR signaling. As activation of MAPK signaling has not been studied in depth in the follicle epithelium we also included a constitutively active Ras allele (Ras85D^V12^, referred to as Ras^V12^) ^31^ to examine the effects of specifically activating the Ras/MAPK cascade. Further, we used constitutively active alleles to overactivate Wnt, JNK and Hippo signaling (arm^S10^, hep^CA^ and yki^S111A.S168A.S250A^, referred to as yki^3SA^) ^32–34^ (Fig. S1A-C). In this study, we focus on the effects of overactive Hh and EGFR-Ras signaling. Overactivation of Hh, EGFR and Ras signaling caused excess follicle cells and were associated with distinct morphological phenotypes (Fig. 1B-E). In ovarioles with Hh^OE^, large clusters of follicle cells accumulated between follicles and the follicles rarely progressed beyond the early stages of oogenesis (Fig. 1C). In ovarioles with overexpression of Ras^V12^, the normal structure of the ovariole was completely disrupted and follicles rarely formed at all (Fig. 1D). Overexpression of λtop^4.4^ also resulted in a large increase in of follicle cells, but they were more evenly distributed on and in between follicles, and most ovarioles contained several mid- and late-stage follicles (Fig. 1E).

We generated single-cell RNA-sequencing data for each genotype in duplicate with 109-30-Gal4 crossed to *w^1118^* as a wildtype control (4 wildtype datasets in total), resulting in a total of 69,994 cells after applying standard quality control filters (Fig S1D-F, Table S1). We merged and batch corrected the data, performed unsupervised clustering, and determined cluster identity in the wildtype dataset using previously described markers (Fig. 1F, S1G, S2A). The clustering agreed well with previously published cell atlases ^12,35–38^, and the increased resolution that we gained by sequencing a larger number of cells provided several new insights. First, we were able to detect two distinct clusters of stalk cells rather than one, as previously reported ^12,35,36^. Both clusters expressed high levels of the stalk cell markers, CG46339 and anterior open (aop) but the cells in one cluster had significantly higher expression of LamC than the other (Fig S2A, Table S2). Staining of wildtype ovaries showed that aop is expressed from Region 3 in the germarium and is thus expressed from the earliest stage of stalk cell differentiation (Fig. S2B) whereas LamC expression is weaker in the first stalk and stronger in later stalk cells (Fig. S2C). Thus, the two stalk cell clusters represent a younger and more mature stalk cell population. Second, we identified a new marker, *biniou* (*bin*) that is strongly enriched in the EC and FSC clusters and very low or undetectable in all other clusters (Fig. S2A). We found that *bin* expression is strongly enriched in FSCs but not pFCs in our wildtype ovary atlas dataset (Fig. S2D). To test this predicted expression pattern *in vivo*, we performed immunofluorescence staining of bin and Fas3 in wildtype ovarioles. Indeed, we observed strong expression in escort cells throughout Regions 1 and 2a, as well as in an average of 3.9 ± 0.57 (n = 15) Fas3^+^ cells at the anterior edge of the Fas3 expression domain, where the FSCs reside (Fig. S2E). These anterior-most Fas3^+^ cells were placed in Layer 1, as defined in ^10^, in 43% of cases and in Layer 2 in the remaining 57%. Third, we found that *deadpan* (*dpn*) is strongly enriched in the polar cell cluster and we confirmed this *in vivo* by immunofluorescence (Fig. S2A, F). Dpn antibody staining reliably identifies polar cells in early cyst stages (Fig. S2F). Lastly, one cluster co-expressed stalk and polar cell markers (Fig. S2A, Table S3). Immunostaining with the stalk cell marker aop and the polar cell marker dpn revealed that posterior polar cells in Stages 2 to 4 occasionally co-express stalk and polar cell markers (Fig. S2F-G).

Three additional clusters were primarily composed of cells from non-wildtype datasets and had expression profiles that are distinct from previously identified wildtype cell types. One cluster which we named “JNK high cells,” was mainly composed of cells from the Ras^V12^, λtop^4.4^ and hep^CA^ datasets and expressed high levels of JNK signaling targets such as *puc*, *Hsp68*, *rpr* and *cher* (Fig. 1F, G, S1G, Table S3) ^39–43^. A second cluster, which we refer to as “DNA damage cells,” primarily contained cells from the Hh^OE^, Ras^V12^ and λtop^4.4^ datasets and expressed DNA damage response genes, including *Irbp*, *DNAlig3*, *Xrp1*, *CG3448* and *Ku80* (Fig. 1F,G, Table S3) ^44–47^. Lastly, one cluster was almost exclusively composed of cells from the Hh^OE^ dataset and its expression profile resembled that of mature stalk cells (Fig. 1F,H, Table S3). We therefore named this cluster “Hh stalk-like”. Together, cluster annotation allowed us to identify cell type specific differentially expressed genes downstream of overactivation of the pro-proliferative signaling pathways Hh, EGFR-Ras, Wnt, JNK and Hippo signaling in follicle cells (Tables S4-9).

### Ectopic Hh signaling induces a transcriptional hybrid state

To better understand the effects that overactivation of Hh, EGFR or Ras signaling has on follicle cell proliferation and differentiation, we quantified the percentage of cells in each mutant genotype (expressed as log_2_-fold change vs wildtype) that contributed to each of the main clusters in the merged dataset (Fig. 2A). In the Hh^OE^ dataset, we observed a substantial increase in the fraction of cells that were assigned to all of the early follicle cell type clusters, including the FSC, pFC, young and mature stalk cell, and polar cell clusters and a decrease in the fraction of Hh ^OE^ cells assigned to all of the MB follicle cell types that are specific for Stage 3 and later. In addition, we observed a two-fold increase in the number of bin^+^, Fas3^+^ cells (8.2 ± 1.08 cells, n = 14, p = 0.0022), which is consistent with our finding that an increased percentage of Hh ^OE^ cells were assigned to the FSC cluster. Differential abundance testing with milo ^48^ confirmed this enrichment of early follicle cell types (Fig. 2B). Further, milo analysis identified overlaps between stalk cell neighborhoods with other cell types (Fig. 2C). We applied multimodal reference mapping to the wildtype ovarian atlas ^12^ to investigate cluster identity in more depth. Mapping of the wildtype dataset reproduced cell identities similar to clusters identified by unsupervised clustering analysis and the cells mapped to these clusters with high confidence (Fig. 1F, 2D-F). In contrast, reference mapping of the Hh^OE^ dataset assigned a large portion of early MB cells to a pFC or stalk cell identity with moderate to low mapping scores (Fig. 1F, 2D-F). To better understand cell type identity upon Hh signaling overactivation, we investigated the expression of known cell type specific markers. We found that *cas*, *zfh1* and *CG46339*, which are highly expressed in wildtype stalk cells, were expressed more broadly throughout the MB follicle populations in Hh^OE^ ovaries (Fig. 2G-M). Moreover, the stalk and polar cells in Hh^OE^ ovaries failed to downregulate the expression of *eya* (Fig. 2G, K). In contrast, expression of aop, which is highly expressed in wildtype stalk cells, was expressed in low levels in early stages of the follicle lineage in Hh^OE^ ovaries and much higher levels in a large set of late stage follicle cells (Fig. 2M). Likewise, LamC and dpn expression were delayed in the stalk and polar cells of Hh^OE^ ovaries (Fig. S3A-D). At later stages, we detected cells co-expressing LamC and eya, which are mutually exclusive in the wildtype ^21^ and, consistent with this, a large set of cells with MB morphology in Hh^OE^ ovaries expressed LamC (Fig. S3E-F).

**Figure 2:**
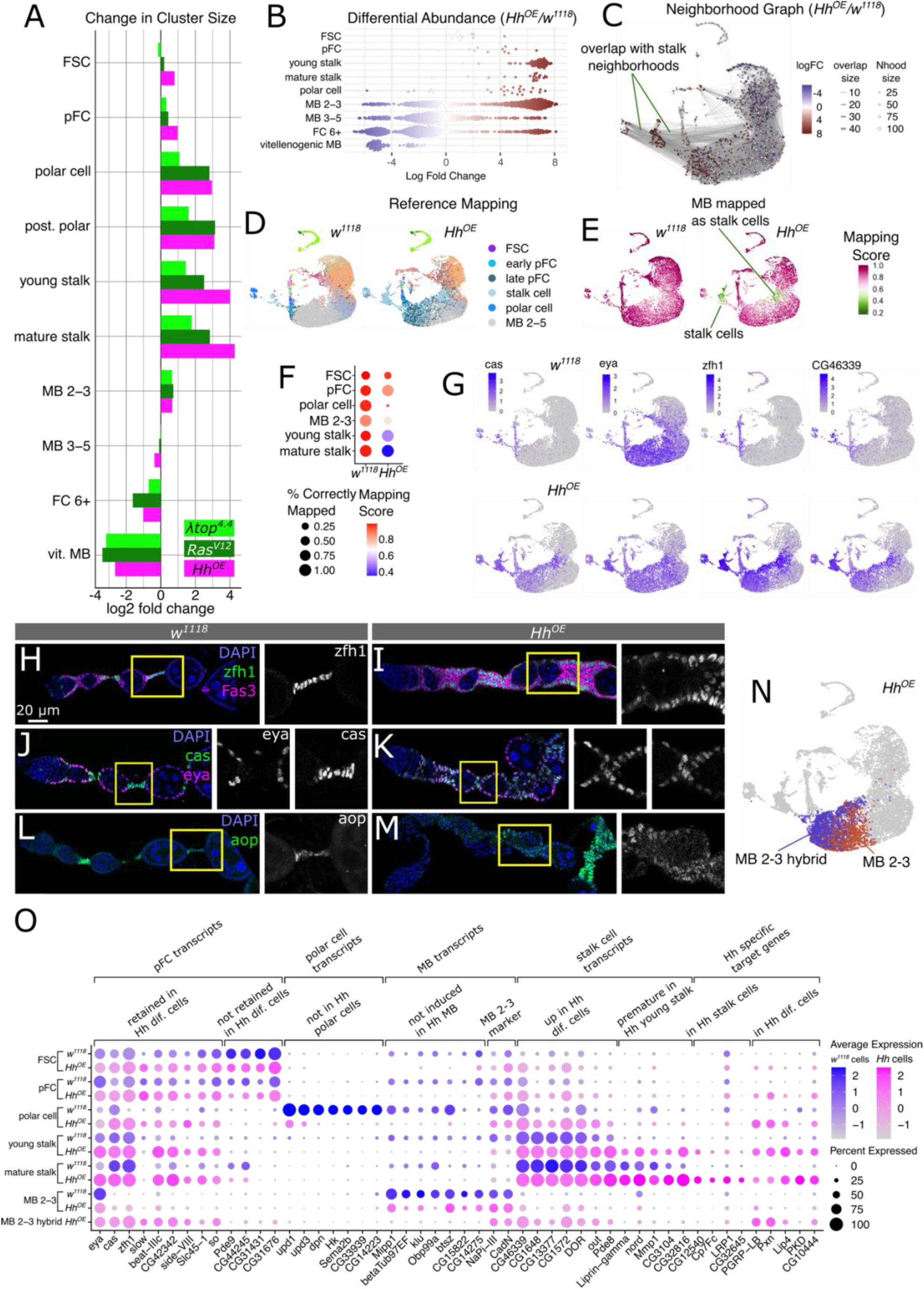
Hh overactivation induces hybrid states in differentiated follicle cells. A) Bar chart displaying the log2 fold change of cluster contribution per cell type and genotype normalized to the respective wildtype population. B) Differential abundance testing of Hh ^OE^ cells in comparison to control *w^1118^* cells. C) A neighborhood graph of the *w^1118^* and Hh^OE^ datasets. Note the multiple neighborhood overlaps between stalk cell neighborhoods and neighborhoods of polar cells and early MB cells. D) UMAP plot of *w^1118^*and Hh^OE^ cells colored upon reference mapping to the *Drosophila* ovary atlas ^12^. Colors of cell types targeted by *109-30*-Gal4 are displayed in the legend. For a full legend refer to Fig. S2H. E-F) Mapping score associated with (D) on a UMAP plot (E) and visualized via a DotPlot (F). G) UMAP plots of *w^1118^* and Hh^OE^ datasets showing the expression of *cas*, *eya* and *zfh1* as indicated. H-M) Ovarioles of *109-30*-Gal4 crossed with *w^1118^* as a control (H,J,L) or UAS-*Hh* (I,K,M) stained for (H-I) DAPI (blue), zfh1 (green) and Fas3 (magenta), (J-K) DAPI (blue), cas (green) and eya (magenta) and (L-M) DAPI (blue) and aop (green). N) UMAP plot of *Hh^OE^* cells displaying sub-clusters of the MB 2-3 cluster. O) DotPlot showing average and percent expression in *w^1118^* and Hh^OE^ clusters containing cells targeted by *109-30*-Gal4.

We noted that only a subset of MB cells in Stages 2-3 upregulated non-MB transcripts (Fig. 2G). Sub-clustering of MB 2-3 produced two clusters out of which one displayed a more wildtype-like expression pattern (Hh^OE^ MB 2-3), while the other cluster displayed an upregulation of Hh target genes (Hh^OE^ MB 2-3 hybrid) (Fig. 2N-O, Table S4). Differential expression analysis of Hh^OE^ clusters targeted by 109-30-Gal4 identified a distinct set of Hh target genes (Fig. 2O, S3G, Table S4). While FSCs and pFCs contained a relatively small number of differentially expressed genes, differentiated cell types retained the expression of multiple genes that are highly expressed in undifferentiated FSCs and pFCs (Fig. 2O, S3G, Table S4). Polar cells and MB cells exhibited decreased expression of cell type specific markers, while simultaneously upregulating stalk cell specific transcripts (Fig. 2O, Table S4). Lastly, young stalk cells prematurely expressed markers of mature stalk cells and all differentiated cells in the 109-30-Gal4 expression domain expressed a set of genes not detected in wildtype follicle cells, which was reflected in cluster tree analysis (Fig. 2O, S3H, Table S4). In summary, these findings show that FSCs and pFCs are largely protected from effects of Hh overactivation, while differentiated cell types retain a portion of an undifferentiated cell expression profile and upregulate stalk cell specific transcripts. This results in a subset of cells acquiring a hybrid state characterized by the expression of genes associated with the undifferentiated state, the stalk cell identity and the polar or MB cell identity.

### Ras^V12^ promotes the polar cell fate

EGFR signaling acts upstream of the Ras pathway in the follicle cells ^16^. However, while EGFR overactivation was extensively studied, activation of Ras has not been investigated in depth. Overactivation of either λtop^4.4^ or Ras^V12^ resulted in an increase of the fraction of cells that were assigned to the polar, stalk, and pFC clusters and a decrease in the fraction of cells assigned to later stages (Fig. 2A, 3A-B). Yet, while expression of λtop^4.4^ had a consistent effect on all cell populations, expression of Ras^V12^ disproportionately increased the polar cell population (Fig. 2A, 3A-B). In addition, reference mapping correctly assigned the majority of λtop^4.4^ cells, but identified a large portion of Ras^V12^ MB cells as polar cells (Fig. 3E-I). Polar cell specific induction of λtop^4.4^ using upd-Gal4 did not result in any morphological defects when compared to a wildtype control (Fig. 3J-K). In contrast, polar cell specific Ras^V12^ expression significantly increased the number of polar cells per cluster and caused a double row stalk phenotype (Fig. 3L-M). Indeed, while λtop^4.4^ and Ras^V12^ shared common target genes, we found that Ras^V12^ specifically targeted a set of genes not induced by λtop^4.4^, including *PKA-C3*, *Arc1* and *CG17839* which are highly expressed in wildtype polar cells (Fig. S4A, Tables S5-6). In addition, we noted a downregulation of the polar cell markers *dpn*, *upd1* and *upd3* in the Ras^V12^ dataset, indicating that Ras^V12^ polar cells do not possess mature polar cell character (Fig. 3N, S3A, Table S6). Consistent with this, immunostaining of ovarioles with pan-follicle cell expression of Ras^V12^ showed a strongly decreased number of dpn^+^ cells, while expression of λtop^4.4^ displayed an increase in polar cells as expected by differential abundance testing (Fig. S3B-C). Interestingly, *LamC* and *cas*, which are highly expressed in stalk cells, were expressed at high levels in the MB cells of the λtop^4.4^ and Ras^V12^ ovaries while stalk but not polar cell populations failed to downregulate *eya* (Fig. 3N-O, Tables S5-6). Consistent with this, we detected cas^+^, eya^+^ and LamC^+^, eya^+^ cells between follicles, where stalk cells reside, in both genotypes (Fig. 3P-S). Together, this indicates that while EGFR and Ras signaling both increase cell number in the early stages of follicle cell differentiation, Ras signaling has an additional role in promoting polar cell differentiation.

**Figure 3:**
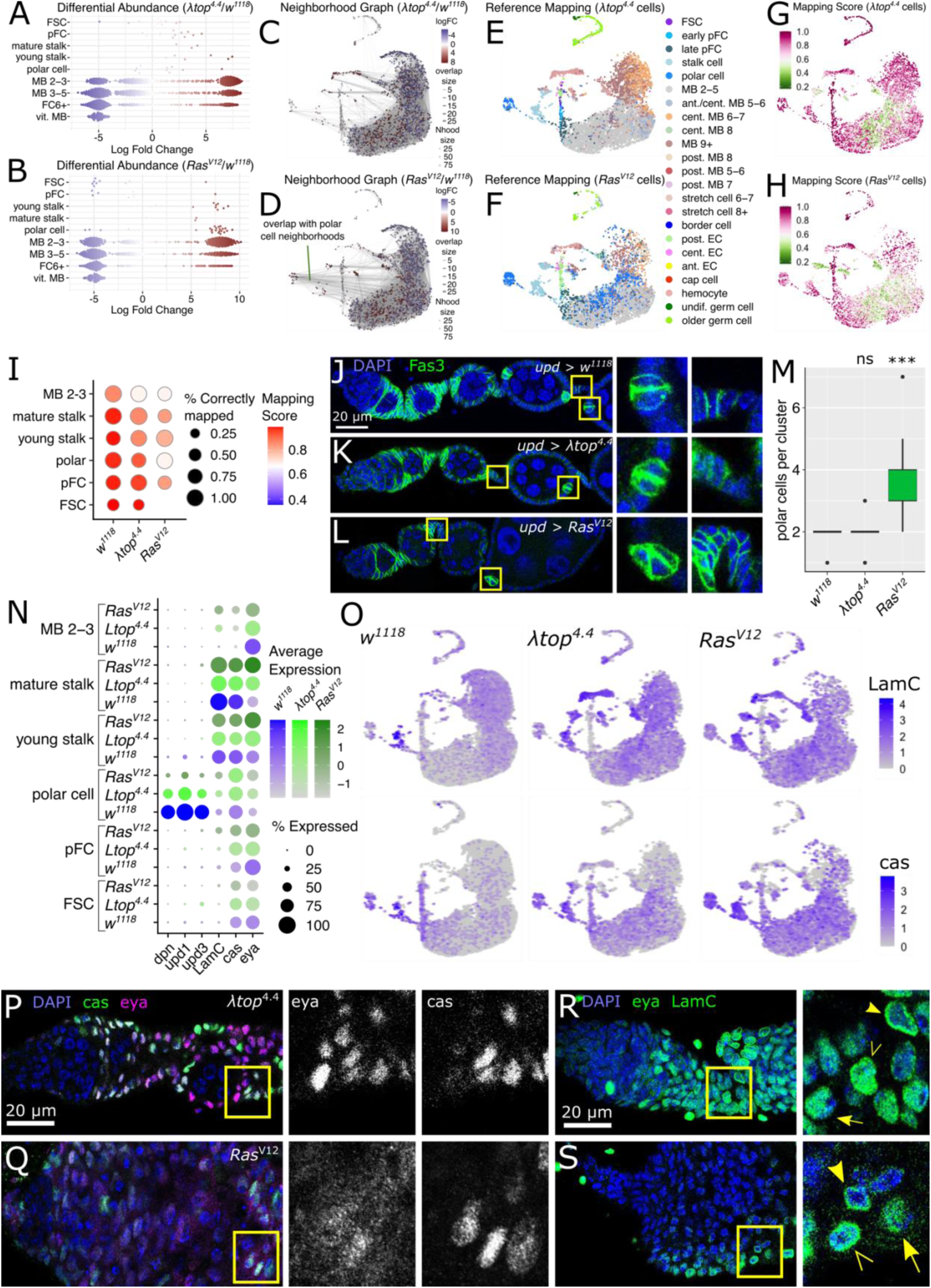
Ras^V12^ but not λtop^4.4^ induces the polar cell fate. A-B) Differential abundance testing of *w^1118^* cells in comparison to (A) λtop^4.4^ or (B) Ras^V12^ cells. C-D) Neighborhood graphs of *w^1118^*cells and (C) λtop^4.4^ or (D) Ras^V12^ cells. Note the multiple overlaps between polar cell and MB neighborhoods in (D). E-F) UMAP plots colored by cluster identity predicted by reference mapping to the *Drosophila* ovary atlas ^12^ of (E) λtop^4.4^ or (F) Ras^V12^ cells. A large number of cells identified as MB by unsupervised clustering are mapped as polar cells in the Ras^V12^ dataset. G-I) Mapping score displayed on a UMAP plot for (G) λtop^4.4^ and (H) Ras^V12^ cells and on a DotPlot (I). DotPlot scale is similar to Fig. 2F to allow for better comparison. J-L) Ovarioles driving Gal4 in polar cells under the *upd* promoter and combined with (J) *w^1118^*, (K) λtop^4.4^ or (L) Ras^V12^ and stained for DAPI (blue) and Fas3 (green). Insets show a mature polar cell cluster (left) and a mature stalk (right). Note that polar cell specific expression of *Ras^V12^* but not *λtop^4.4^* or the control results in more than 2 polar cells per cluster and double row stalks. M) Boxplot showing the quantification of polar cells per cluster (St. 5-7) of genotypes in (J-L). n = 25, 23, 18 ovarioles for *w^1118^*, λtop^4.4^ and Ras^V12^ respectively. p from Kruskal-Wallis test for λtop^4.4^ = 0.9485 (ns), Ras^V12^ = 1.791e-08 (***). N) DotPlot showing the expression of the polar cell marker *dpn*, the stalk cell marker *LamC* and *cas* and *eya* in the *w^1118^*, λtop^4.4^ and Ras^V12^ clusters with *109-30*-Gal4 activity. O) UMAP plots showing the expression of *LamC* and *cas* in the *w^1118^*, λtop^4.4^ and Ras^V12^ datasets as indicated. Note that a subset of MB cells in the *λtop^4.4^* and *Ras^V12^* expresses *LamC* and *cas.* P-S) Ovarioles expressing λtop^4.4^ (P,R) or Ras^V12^ (Q,S) under the *109-30*-Gal4 driver and stained for DAPI (blue) and cas (green) and eya (magenta) (P-Q) or eya and LamC (green (R-S). Insets show cells beyond the FSC/pFC domain which co-express cas and eya (P-Q) or eya and LamC (R-S). Arrow points out eya^+^ cells, filled arrowheads highlight LamC^+^ cells and empty arrowheads points at eya^+^ LamC^+^ cells.

### Aberrant EGFR-Ras signaling represses main body follicle cell differentiation

Specific transcription factor activities are the main drivers of cell fate and behavior. We sought to identify transcription factors with differential activity in our datasets. To this end we applied single - cell regulatory network interference and clustering (SCENIC) ^49^. SCENIC examines the expression of a transcription factor and identifies co-expressed genes as potential transcription factor targets. This conjunction of transcription factor and target genes constitute a “regulon”. When we identified regulons for each genotype, we noted discrepancies between the sets of genes contributing to the regulons. To allow for direct comparison of transcription factor activity between genotypes, we focused on regulons generated by SCENIC analysis of the wildtype datasets and tested for significant differences between wildtype and mutant genotypes (Fig. S5A, Tables S10-11). The regulons induced by pointed (pnt) and E2F transcription factor 1 (E2f1) and the E2f1 interactor Dp, all of which are known to regulate proliferation ^50,51^, were increased in the λtop^4.4^, Ras^V12^ and Hh^OE^ datasets (Fig. 4A, S5A). This is in line with increased proliferation induced by λtop^4.4^, Ras^V12^ and Hh^OE^ ectopic expression. Further, we detected activity of zfh1 and Mef2 particularly in Hh^OE^ hybrid cells, while in the *w^1118^* dataset activity of both regulons was largely restricted to stalk cells (Fig. 4A, S5A). Previous studies have identified zfh1 as a Hh target gene in the *Drosophila* testis ^52,53^ and we detected increased *zfh1* RNA levels downstream of Hh signaling (Fig. 2G). This suggests that zfh1 is induced by Hh signaling in follicle cells to promote the stalk cell fate. In contrast, Hh overactivation did not induce *Mef2* RNA expression (Fig. S5B), indicating that Hh induces Mef2 activity indirectly. Hh signaling as well as the mammalian orthologs of zfh1, ZEB1 and 2, and Mef2 have been implicated in epithelial-to-mesenchymal transition (EMT) ^54–56^. To test whether Hh signaling induces EMT in follicle cells we investigated the expression of other regulators of EMT. Indeed, Hh signaling induced the expression of *pebble* (*pbl*) and *heartless* (*htl*), two markers of mesenchymal fate (Fig. S5B) ^57,58^.

**Figure 4:**
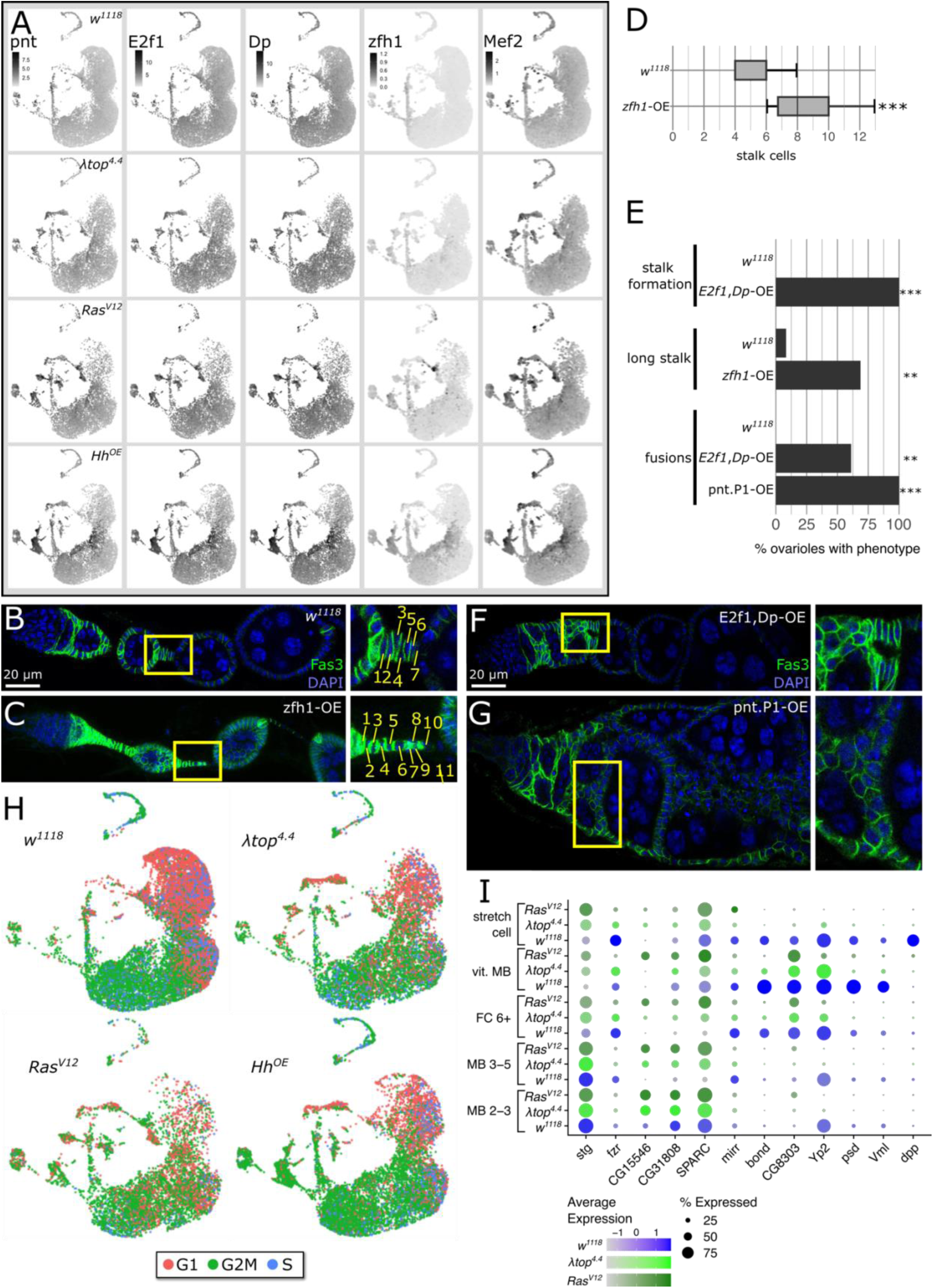
EGFR-Ras signaling regulates main body follicle cell differentiation. A) UMAP plots of the indicated genotypes showing the regulon activity of the indicated transcription factor. High confidence regulons were identified by SCENIC ^49^ analysis of the *w^1118^* wildtype control datasets and expression analyzed using PercentageFeatureSet() of the Seurat package ^89^. B-C) Ovarioles of *109-30^ts^*crossed with the wildtype *w^1118^* (B) or inducing overexpression of zfh1 (C) stained for DAPI (blue) and Fas3 (green). Insets show a mature stalk region with individual stalk cells numbered. D) contains the associated quantification of cells per mature, single-row stalk. n = 13 and 16 stalks for *w^1118^* and zfh1-OE respectively. p from student’s t test = 7.59e-05. E) Barplot presenting the penetrance of phenotypes induced by transcription factor overexpression. n: stalk formation: *w^1118^* = 13, E2f1,Dp-OE = 18, long stalk: *w^1118^* = 12, zfh1-OE = 16, fusions: *w^1118^* = 13, E2f1-Dp-OE = 18, pnt.P1-OE = 18 ovarioles. p values from student’s t-test: stalk formation: E2f1,Dp-OE = 2.00e-07, long stalk: zfh1-OE = 0.00281, fusions: E2f1-Dp-OE = 0.001755, pnt.P1-OE = 2.00e-07. F-G) Ovarioles of *109-30^ts^* inducing overexpression of E2f1 and Dp (F) or pnt (G) and immunostained for DAPI (blue) and Fas3 (green). Insets show regions with morphological abnormalities in. H) UMAP plot of *w^1118^*, λtop^4.4^, Ras^V12^ and Hh^OE^ cells scored for cell cycle phase. Note the switch from predominantly G2M to predominantly G1 phase in *w^1118^*and Hh^OE^ MB cells, while no clear switch is visible in the λtop^4*.4*^ and Ras^V12^ datasets. I) DotPlot of MB clusters of *w^1118^*, λtop^4.4^ and Ras^V12^ datasets showing average and percent expression of genes inducing the mitosis to endocycle switch (*stg*, *fzr*) and stage specific MB markers. Markers of early MB Stages (*CG15546*, *CG31808*, *SPARC*) are not downregulated in lates stages of MB fates in the λtop^4.4^ and Ras^V12^ datasets, while markers of differentiated MB cell fates are not induced (*mirr*, *bond*, *CG8303*, *Yp2*, *psd*, *Vml*, *dpp*).

We performed overexpression targeting these transcription factors to confirm that a misregulation is functionally relevant. Overexpression of zfh1 resulted in long stalks with a significantly increased number of cells per stalk when compared to a wildtype control (Fig. 4 B-E), confirming zfh1 as one of the Hh targets inducing stalk cell fate. In addition, overexpression of E2f1 and Dp or pnt resulted in prominent fusions of cysts resembling the λtop^4.4^ overexpression phenotype (Fig. 1E, 4E-G). E2f1 is a well-known cell cycle regulator required for G1 to S transition and pnt induces cell cycle related genes during neural development and is induced by Ras signaling ^50,51,59^. This strongly suggests that increased proliferation is a major driver of the phenotype observed upon λtop^4.4^ overexpression. To test this hypothesis further, we examined the expression of known cell cycle specific genes in our dataset to determine the cell cycle phase for each cell (Fig. 4H, S5C, Table S12). We found that MB follicle cells have a long G2 phase during mitotic divisions up to Stage 5, and switch to a long G1 phase from Stage 6 onwards, at which point MB cells start endoycling (Fig. 4H, S5C) ^60^. While overactivation of Hh signaling had a negligible effect on the distribution of cell cycle phases, EGFR and Ras signaling delayed the G2 to G1 switch in MB follicle cells (Fig. 4H, S5C). We examined the expression of genes known to regulate the switch to endocycling, including *string* (*stg*), which is downregulated, and *fizzy-related* (*fzr*), which is upregulated to facilitate the exit from mitotic cell cycle ^61^. While the *w^1118^*control dataset displayed the expected expression changes of *stg* and *fzr* in MB differentiation, MB populations from the λtop^4.4^ or Ras^V12^ datasets maintained *stg* expression and failed to upregulate *fzr* in later MB fates (Fig. 4I), suggesting the ectopic EGFR-Ras signaling blocks endocycling. Studies on embryonic stem cells showed that a long G1 phase renders cells susceptible to differentiation ^62–64^. We hypothesized that the reduction of late stage MB follicle cell populations observed in the λtop^4.4^ or Ras^V12^ datasets (Fig. 2A, 3A-B) is caused by a failure of MB cells to differentiate properly. Indeed, MB populations in the λtop^4.4^ or Ras^V12^ datasets retain markers for early MB populations like *CG15546*, *CG31808* and *SPARC* and fail to upregulate markers of MB differentiation including *mirr*, *bond*, *CG8303*, *Yp2*, *psd*, *Vml* and *dpp* (Fig. 4I) ^12,35,36^. These data suggest that sustained EGFR-Ras signaling blocks the entry of MB follicle cells to the endocycle and that the switch to endocycling is crucial for proper MB differentiation.

### Tight control of independent signaling pathways prevents malignant cell growth

Interestingly, among all datasets we noted comparably mild changes in gene expression in the FSC and pFC clusters (Fig. 2O, 3N, S6A, Tables S4-9). Likewise, we observed only small increases in the number of cells in the FSC and pFC populations of these mutant ovarioles (Fig. 2A). To investigate the impact on FSC number, we quantified the frequency of *bin*^+^, *Fas3^+^* double positive cells in follicle cells with signaling pathway overactivation. Expression of arm^S10^ and Hh^OE^ resulted in a roughly 2-fold increase in stem cell number (Fig. 5A). These results suggest that multiple signaling pathways may work together to confer robustness that protects the tissue from overexpansion of the stem cell pool. To test this hypothesis, we co-overactivated pairs of signaling pathways and examined changes in FSC number. First, we overactivated pairs of signaling pathways that have previously been described to interact in the stem cells. Wnt signaling acts upstream of EGFR-Ras signaling in FSCs ^65^. Consistent with this, concurrent overexpression of arm^S10^ and λtop^4.4^ together did not increase FSC number compared to arm^S10^ overexpression alone (Fig. 5A). Similarly, co-overactivation of Hh and Hippo signaling, which interact in FSCs ^19,20^, does not expand FSC numbers beyond the increase observed by Hh overexpression alone (Fig. 5A). In contrast, co-overactivation of signaling pathways which are not known to interact in FSCs had a synergistic effect on the increase in FSC number (Hh^OE^ with arm^S10^: 18.8 ± 1.81, λtop^4.4^: 21.1 ± 2.39, Ras^V12^: 62.7 ± 6.31 or hep^CA^: 19,3 ± 2.1) (Fig. 5A). Thus, these pathways operate in a combinatorial manner to promote stem cell self-renewal.

**Figure 5:**
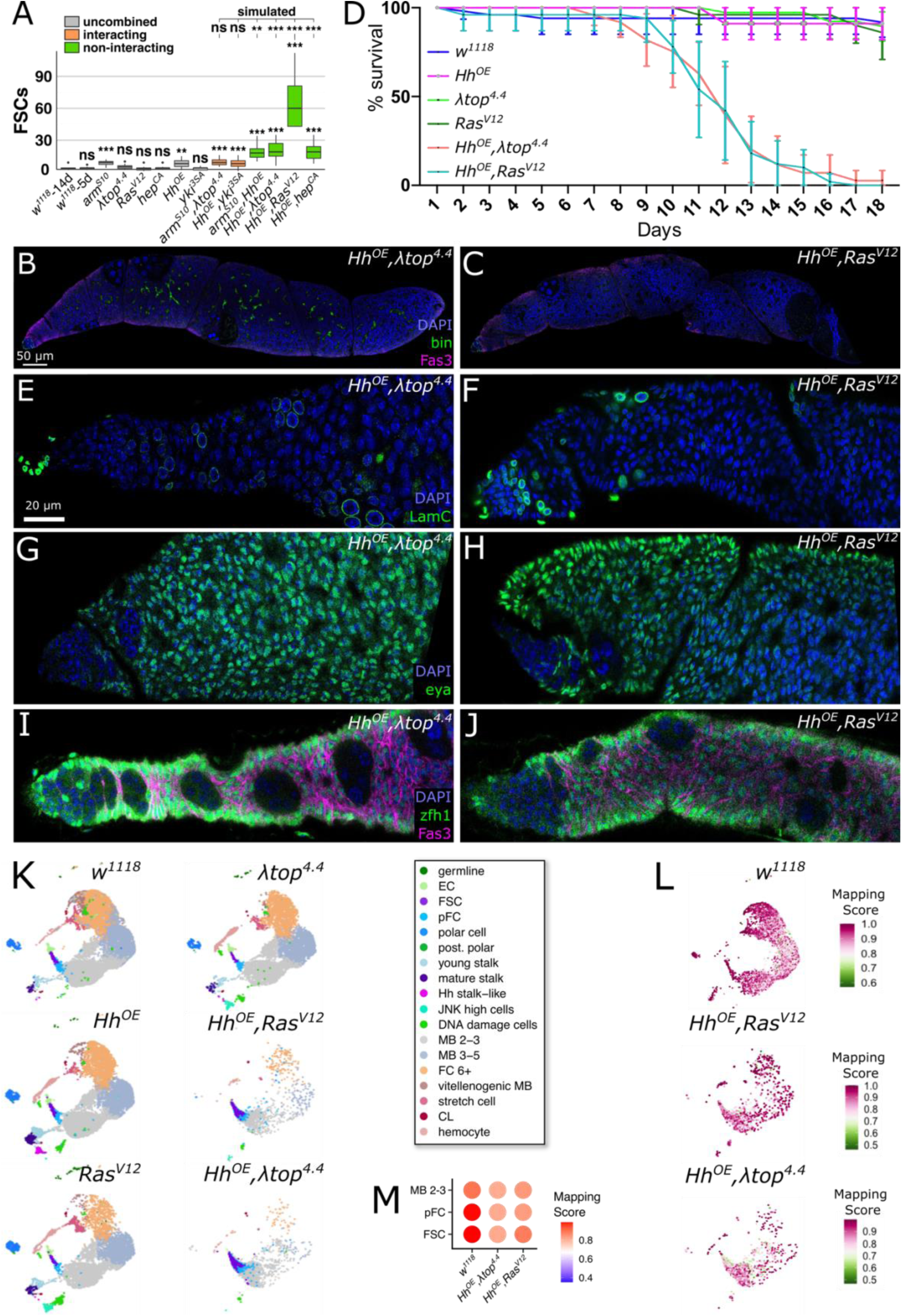
Co-overactivation of Hh and EGFR-Ras pathways induces follicle cell tumors. A) Box chart showing the number of bin^+^Fas3^+^ cells in each of the indicated genotypes induced by *109-30^ts^*. Double overactivations are separated into those interacting and non-interacting in FSCs. Simulated values were calculated by adding the difference of bin^+^Fas3^+^ cell number in each single overactivation compared to the wildtype. n = 5 germaria per genotype. n ≥ 13 germaria for each condition. p-values from student’s t-test defined as p ≥ 0.05: ns, p < 0.05: *, p < 0.01: **, p < 0.001: ***. B-C) Morphology of an entire ovariole stained for DAPI (blue), bin (green) and Fas3 (magenta) expressing (B) Hh^OE^ and λtop^4.4^ or (C) Hh^OE^ and Ras^V12^ under the *109-30* driver. D) Survival assay of flies with *109-30^ts^* driving the indicated genotypes. Flies with follicle cell-specific overexpression of Hh^OE^ and either λtop^4.4^ or Ras^V12^ have a drastically reduced life span. n = 46, 28, 58, 33, 50, 40 flies from 5 different repeats for *w^1118^*; λtop^4.4^; Ras^V12^; Hh^OE^; Hh^OE^,Ras^V12^; Hh^OE^,λtop^4.4^ respectively. p-values from Kaplan Meier analysis defined as p ≥ 0.05: ns, p < 0.05: *, p < 0.01: **, p < 0.001: ***. E-J) Immunostaining of ovarioles of 109-30ts driving Hh^OE^,λtop^4.4^ or Hh^OE^,Ras^V12^ as indicated. E-F) Ovarioles stained for DAPI (blue) and LamC (green). G-H) Ovarioles stained for DAPI (blue) and eya (green). I-J) Ovarioles stained for DAPI (blue), zfh1 (green) and Fas3 (magenta). K) UMAP plots of datasets of the indicated genotypes induced by *109-30^ts^*. Samples of dataset 2 were reference mapped to dataset 1. Note that reference mapping required us to re-run the calculation of UMAP coordinates. Cluster identities of Ras^V12^, λtop^4.4^ and Hh^OE^ datasets are equal to those reported above. L-M) Mapping score for the reference mapping of the Hh^OE^,Ras^V12^ and Hh^OE^,λtop^4.4^ datasets on the UMAP (L) and visualized via a DotPlot (M). The scale of the DotPlot is similar to those reported above to allow comparison.

The co-overexpression of two signaling pathways resulted in strong morphological defects, as expected (Fig. 5B-C, S6B-E), and the combination of Hh^OE^ with either λtop^4.4^ or Ras^V12^ produced exceptionally large tumors. Interestingly, this was associated with a significantly reduced lifespan, with fewer than 10% of mutant flies surviving past 18 days (Fig. 5D). To further investigate this phenotype, we assessed the impact of Hh and EGFR-Ras activation on several cell type specific markers. The patterns of marker expression that we observed strongly suggest that the cells are arrested at an earlier stage of differentiation than they would be with overexpression of any of the three pathways alone (Fig 5E-J). Specifically, all cells were zfh1^+^ and eya^+^, whereas overexpression of these three pathways alone increased the number of cells that expressed stalk cell markers, overactivation of Hh and EGFR or Hh and MAPK caused a substantial reduction in the number of aop^+^ and LamC^+^ cells (Fig 5E-J). Hence, overactivation of Hh with overactivation of either EGFR or Ras causes both an expansion of the FSC pool and maintains downstream follicle cells in an undifferentiated pFC-like state. To confirm this, we performed single-cell RNA-sequencing of these Hh-EGFR/Ras follicle tumor cells along with a paired *w^1118^*control (dataset 2). We sequenced a total of 20,308 high quality cells, which we reference mapped to the dataset containing respective control genotypes (*w^1118^*, Hh^OE^, λtop^4.4^ or Ras^V12^, dataset 1). Reference mapping successfully mapped both additional *w^1118^* datasets resulting in an integrated control containing all six *w^1118^* datasets (Fig 5K-M, S6F). The hereby identified wildtype clusters of the two additional *w^1118^* datasets further recapitulated the expected expression pattern of cell type specific markers and contained cells of all cluster identities (Fig. S6G, Table S13). In contrast, cells from the datasets with Hh^OE^,λtop^4.4^ or Hh^OE^,Ras^V12^ primarily contained cells with FSC, pFC and MB Stage 2-3 identities although they were mapped with relatively low confidence (Fig. 5K-M, Table S13). This result was consistent among both Hh^OE^,Ras^V12^ datasets and the Hh^OE^,λtop^4.4^ dataset (Hh^OE^,λtop^4.4^-1), as well as a second Hh^OE^,λtop^4.4^-2 dataset, which we excluded from further analysis due to low data quality (Fig. S6F, H-J). Indeed, when we examined the percentage of cells contributing to each cluster, we found a massive increase in the contribution to the FSC and pFC clusters and a slight increase in early MB follicle cells (MB 2-3) in both genotypes (Fig. 6A, Table S13). Differential abundance testing of the Hh^OE^,Ras^V12^ dataset confirmed that there was a substantial increase of undifferentiated cell types and a decrease or complete absence of differentiated cell types (Fig. 6B). Moreover, neighborhood analysis identified a high amount of overlap between the remaining neighborhoods containing differentiated follicle cells with pFC neighborhoods (Fig. 6C). These data provide confirmation that Hh-EGFR/Ras follicle tumor cells largely retain a pFC character. To test which cell type gives rise to the tumor, we made use of Gal4 driver lines targeting distinct follicle cell populations to express Hh^OE^ with either λtop^4.4^ or Ras^V12^ ^12^. While the induction in FSCs using the Wnt4-Gal4 driver, or in stalk cells using the CG46339-Gal4 driver resulted in morphological phenotypes, only induction with the stl-Gal4 driver, which targets follicle cells from the pFC fate onwards, resulted in a tumor-like growth (Fig. 6D-I) ^12^. This demonstrates that these Hh-EGFR/Ras follicle tumor cells not only retain pFC character, but the tumor is also initiated from cells with pFC fate.

**Figure 6:**
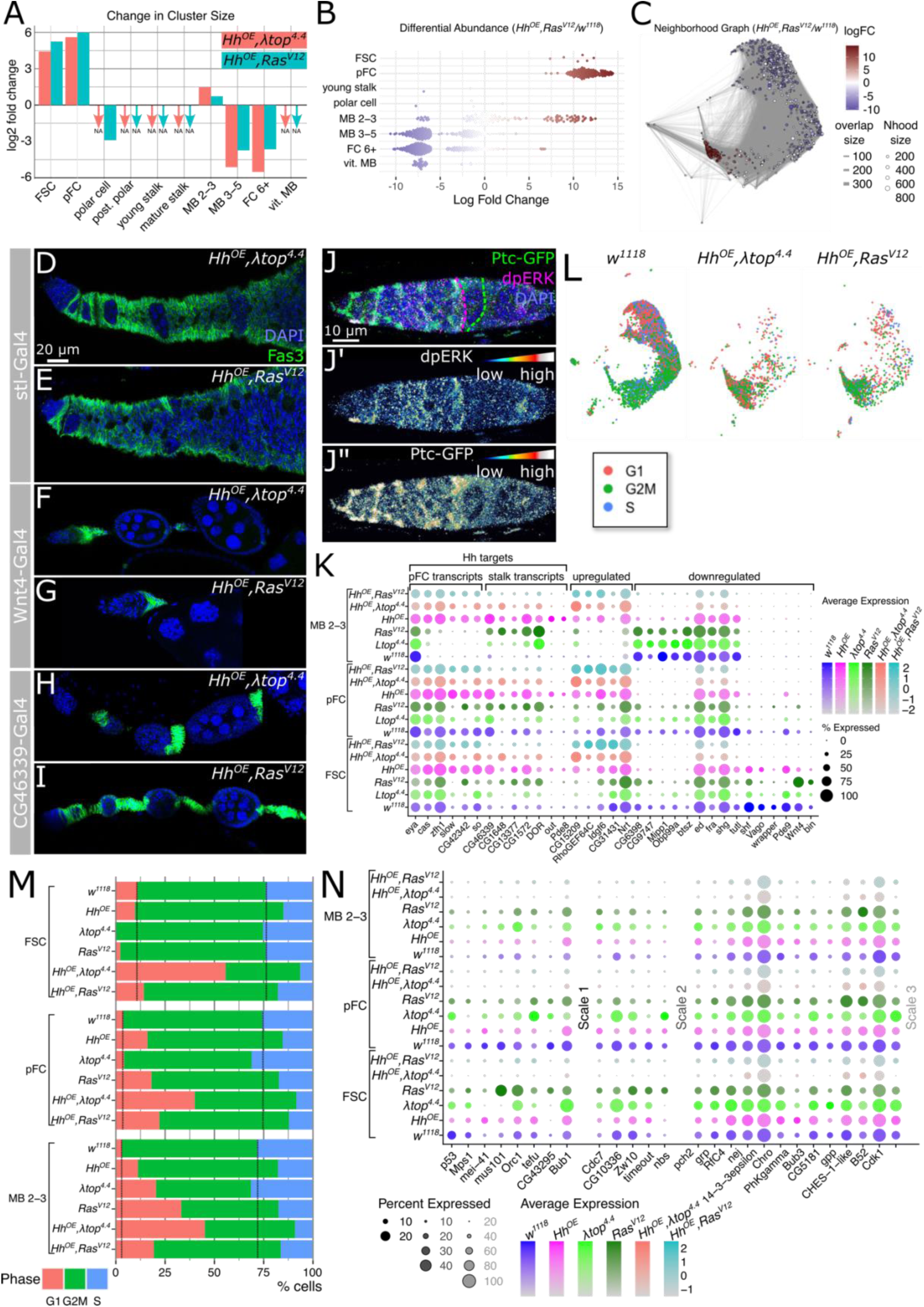
Hh-EGFR/Ras co-overactivation prevents differentiation and cell cycle checkpoints. A) Bar chart showing the log2 fold change of cluster contribution per cell type for the Hh^OE^,Ras^V12^ and Hh^OE^,λtop^4.4^ datasets normalized to the respective wildtype population of the paired *w^1118^* control datasets. NA marks absent cell populations. B) Differential abundance testing identifies a strong increase in FSCs and pFCs and a decrease of differentiated follicle cell types in the Hh^OE^,Ras^V12^ datasets in comparison to the paired *w^1118^*datasets. C) Graph showing overlap between neighborhoods of the Hh^OE^,Ras^V12^ dataset with the paired *w^1118^* dataset. D-I) Ovarioles with Hh^OE^,λtop^4.4^ or Hh^OE^,Ras^V12^ induced in distinct subsets of follicle cells and stained for DAPI (blue) and Fas3 (green). D-E) Induction with *stl*-Gal4, which is active from pFCs onwards, F-G) with *Wnt4*-Gal4, which is inactive after the FSC state, H-I) *CG46339*-Gal4, a stalk cell driver. Induction with *stl*-Gal4 phenocopies the induction with the pan-follicle cell driver *109-30*-Gal4. J) Wildtype germarium expressing Ptc-GFP (green) and stained for dpERK (magenta) and DAPI (blue). dpERK and Ptc-GFP are shown in colorimetric scale in (J’) and (J”), respectively. dpERK expression boundary is outlined with magenta and corresponds to the FSC/early pFC region. Ptc-GFP expression boundary is outlined with a green dotted line. A small population of cells posterior (right) to the dpERK expression boundary maintains Ptc-GFP expression and corresponds to the region harboring late pFCs. K) DotPlot showing average and percent expression in FSCs, pFCs and MB cells of Stages 2-3. Hh target genes which are expressed in wildtype pFCs are upregulated in the double overactivation datasets, while Hh targets which are stalk specific are not induced. L) UMAP plots of the paired wildtype control *w^1118^*and the Hh^OE^,λtop^4.4^ and Hh^OE^,Ras^V12^ datasets colored for the predicted cell cycle phase. M) Bar graph showing the percentage of cells per cluster in G1, G2M and S phase of the cell cycle. Note the increase of cells in G1 phase in the Hh^OE^,λtop^4.4^ and Hh^OE^,Ras^V12^ datasets. N) DotPlot showing the expression of cell cycle checkpoint regulators, all of which are downregulated in the Hh^OE^,λtop^4.4^ and Hh^OE^,Ras^V12^ datasets, particularly in FSCs and pFCs. Note that the low expression of some of the cell cycle regulators required the use of three scales.

Our analysis showed that Hh induces pFC and stalk cell transcripts (Fig. 2, Table S4). However, Hh overexpression did not induce stalk cell specific transcripts including *CG1648*, *out* and *Pde8* in the FSC and pFC populations at high levels (Fig. 2O). Taken together, these results suggest that Hh signaling does not induce stalk cell transcripts in cells with active EGFR-Ras signaling. Consistent with this, we confirmed that EGFR-Ras activity decreases well before Hh signaling activity, as indicated by immunostaining for dpERK and the ptc-GFP reporter (Fig. 6J). In addition, we found that simultaneous overactivation of the Hh and EGFR-Ras pathways induced high levels of pFC-specific Hh targets, but not stalk cell specific genes induced by Hh signaling (Fig. 6K, Tables S14-15). Moreover, we identified a substantial number of differentially expressed genes that encode regulators of basic epithelial properties, like cell adhesion (*Nrt*, *btsz, ed, fra, shg, tutl, wrapper*), cell polarity (*RhoGEF64C*) or production of the extracellular matrix (*Idgf6*) (Fig. 6K, Tables S14-15). In particular, the downregulation of cell adhesion molecules like E-Cadherin, encoded by *shg* in *Drosophila*, is a hallmark of cancer cells and is likely the cause for the multilayering of Hh-EGFR/Ras follicle tumor cells (Fig. 5B-C) ^66^. Taken together, these findings support the hypothesis that Hh signaling promotes the undifferentiated FSC/pFC state in cells with active EGFR-Ras signaling, while promoting differentiation towards the stalk cell fate upon a decline of EGFR-Ras activity.

The massive increase of follicle cells in the Hh-EGFR/Ras tumors and the misregulation of the cell cycle we observed by ectopic expression of λtop^4.4^ and Ras^V12^ prompted us to investigate the cell cycle of the Hh-EGFR/Ras follicle tumor cells in more depth. Contrary to our expectations, a large proportion of Hh-EGFR/Ras tumor cells of either identity was predicted to be in G1 phase, while cells from both *w^1118^* control datasets behaved like the initial control (Fig. 6L-M, compare to Fig. 4H, S5C). We suspected that these tumor cells pass through the cell cycle more rapidly, causing the observed overproduction of follicle cells. In agreement, a substantial number of genes regulating cell cycle checkpoints, like *p53* and the Chk1 encoding *grapes* (*grp*), were downregulated in Hh-EGFR/Ras tumor cells (Fig. 6N) ^67,68^. Together, our data suggest that while Hh signaling maintains an undifferentiated state in tumor cells, the EGFR-Ras pathway promotes uncontrolled proliferation, together resulting in a massive production of follicle cells. In addition, a misregulation of cell adhesion genes allows the tissue to form dysplasia-like multilayers instead of organizing into a single layer epithelium, like a healthy MB population. Interestingly, many differentially expressed genes in Hh-EGFR/Ras tumor cells are not targeted by the activation of either single pathway (Fig. 6K, N, S7). Together, this indicates that Hh and EGFR-Ras signaling cooperatively induce a set of target genes which confers malignancy.

## Discussion

Here we discovered new points of interaction between signaling pathways that promote self-renewal and differentiation in the FSC lineage. The spatial arrangement of these cues must be tightly controlled to simultaneously allow for correct proliferation rates and specification of differentiation fates within a relatively small region of the tissue. Our findings demonstrate that the interaction between Hh and EGFR-Ras signaling is a key regulator of this process, with Hh signaling toggling from a self-renewal cue to a differentiation cue in the context of low versus high EGFR-Ras signaling, respectively. In wildtype tissue, concurrent activation of both pathways in FSCs is required for self-renewal and, therefore, tissue homeostasis. Yet the overactivation of both EGFR-Ras and Hh throughout the early FSC lineage has a highly detrimental effect leading to the formation of lethal tumors. This highlights the importance of spatially restricting self-renewal signals to the niche and suggests that pathway interactions, such as the EGFR-Ras and Hh interaction that we describe here, may provide failsafe mechanisms that add robustness to the process.

Strikingly, we found that the induction of the Hh transcriptional program results in a novel “hybrid state,” in which cells simultaneously express markers of multiple distinct differentiation states as well as an undifferentiated state. This illustrates that the identification of the differentiation state of the cell requires the simultaneous analysis of multiple cell fate markers. Similar hybrid states occur during cancer, where cells with an epithelial/mesenchymal hybrid state are the main drivers of metastasis ^69^. Interestingly, ZEB1 and 2, the orthologs of zfh1 which we identified as a transcription factor positively regulated by Hh signaling, are master regulators of this epithelial/mesenchymal hybrid state ^55,56,69^. Hh signaling was shown to induce EMT in other systems and here we describe Mef2, *pebble* and *heartless* as novel downstream targets of Hh signaling, all of which have been implicated in EMT ^54,57,58,70,71^. All these EMT-associated factors are highly expressed in wildtype stalk cells which undergo cell intercalation to form a single row of cells ^72,73^. Many forms of collective cell migration are accompanied by partial EMT ^74^, raising the possibility that stalk cell intercalation is supported by partial EMT and induced in early stages of pFC differentiation by Hh signaling.

Overactivation of EGFR or Ras signaling induces the activity of E2f1, Dp and Pointed, in agreement with previous studies ^59,75,76^. Of note, these transcription factors are relevant downstream targets of EGFR/Ras in the context of cancer ^77,78^. While E2f1-Dp, promote G1 to S progression, the human orthologs of Pointed, ETS1 and 2, promote G2 to M progression ^79,80^. EGFR and Ras particularly induce cell cycle defects in MB follicle cells, which undergo a switch from mitosis to endocycling ^61^. EGFR and Ras disrupt this switch via promoting *string* expression and repression of *fizzy-related* ^61^. Here, we find that the mitosis to endocycle switch of MB cells is accompanied by a lengthening of the G1 phase. An extended G1 phase is tightly linked to differentiation ^62–64^ and in agreement MB follicle cells with active EGFR/Ras signaling fail to differentiate into distinct MB populations.

The simultaneous activation of EGFR-Ras and Hh signaling results in tumorous overproliferation of follicle cells. These follicle cell tumors are driven by an expansion of transit amplifying cells. The overactivation of both Hh and EGFR/Ras causes a misregulation of the cell cycle that is similar to the effect of overactivating either EGFR or Ras alone. In addition, these tumor cells induce a number of genes not targeted by either signaling pathway alone, suggesting that Hh and EGFR-Ras act synergistically as shown in certain cancer types ^6^. Among the cooperatively regulated target genes are regulators of various cell cycle checkpoints, like *p53* and the G2 checkpoint kinase gene *Wee1.* A failure to induce cell cycle checkpoints renders cancer cells highly proliferative while circumventing cell death ^67^. Additionally, Hh and EGFR-Ras cooperatively regulate cell adhesion genes, including E-cadherin, the downregulation of which is a likely cause of multilayering in the tissue. The simultaneous activation of EGFR-Ras and Hh signaling further prevents EMT-like hybrid states and markers of differentiation induced by Hh signaling alone. Interestingly, combined treatment targeting EGFR/Ras and Hh signaling has been shown to significantly reduce metastasis ^81,82^. Thus, our findings suggest that one benefit of simultaneously inhibiting both the EGFR-Ras and Hh pathways is that it may prevent the establishment of epithelial/mesenchymal hybrid states that possess high metastatic potential. Furthermore, inhibition of EGFR-Ras in cancer cells with active Hh signaling may in fact be detrimental if it promotes a switch from a highly proliferative state to a state with high metastatic potential. This highlights the need for further research to understand how Hh and EGFR-Ras signaling intersect in cancer development and suggest that a combinatorial targeting of both signaling pathways should be broadly applied.

How Hh and EGFR-Ras signaling interact in tumorigenesis are ongoing questions in cancer research ^83,84^. Our findings not only provide insight into the underlying mechanisms and target genes but also identify a genetically modifiable model that closely resembles several distinct steps of tumor development. In addition, tumor host flies display a reduced lifespan. While the tumor tissue does not exceed normal ovaries in weight, tumor-bearing flies display a reduced fat and muscle tissue (data not shown), signifying a cachectic phenotype. This might allow the use of the Hh-EGFR/Ras tumor model not only for future studies addressing the specific functions of these signaling pathways but also for investigations of cancer cachexia as previously demonstrated by the use of another ovarian tumor model ^85^.

### Limitations of the study

In our study we use the *109-30-Gal4* driver to induce gene expression. While we are not aware of any other genetic manipulations induced with this driver that decrease the lifespan of the fly, we cannot exclude that tumor-host lethality is caused by the expression elsewhere in an essential organ rather than by tumorigenic follicle cells. Further, we apply ectopic expression of alleles resembling oncogenes, which limits our ability to deduce functions of signaling pathways in the wildtype condition.

## Materials and Methods

### Fly husbandry

Flies were reared at 18°C and newly hatched adults were shifted to 29°C and fed wet yeast daily until dissection. Flies were dissected 14 days after the temperature shift if possible. Genotypes that presented as lethal after 14 days, in particular double overactivation of Hh^OE^ and λtop^4.4^ or Hh^OE^ and Ras^V12^, were dissected after 5 days instead.

The following stocks were used:

BDSC: tub-Gal80^ts^ #7108 and #7017, 109-30-Gal4 #7023, Wnt4-Gal4 #67449, stl-Gal4 #77732, CG46339-Gal4 #77710, w^1118^ #3605, UAS-arm^S10^ #4782, UAS-Ras85D^V12^ #64196, UAS-hep^CA^ #6406, UAS-Hh::EGFP #81024, UAS-yki^S111A.S168A.S250A^ #28817, UAS-pnt.P1 #869, UAS-E2f1, UAS-Dp #4770, UAS-zfh1 #6879, UAS-Jra #7216,UAS-kay-RNAi #33379. VDRC: UAS-Jra-RNAi #107997. The following lines were kindly gifted as indicated: Ptc-pelican (T. Kornberg, UCSF, San Francisco), upd1-Gal4 (D. Montell, UCSB, Santa Barbara), UAS-EGFR^λtop4.4^ (T. Schüpbach, Princeton University, Princeton), Fz3-RFP (gift from R. DasGupta, Genome Institute of Singapore, Singapore), AP1::GFP (T. Harris, University of Toronto, Toronto).

### Immunofluorescence staining and imaging

Ovaries of adult flies were dissected at RT in PBS and fixed for 20 min at RT with 4% PFA. The tissue was washed twice for 10 min in PBS and blocked for 30 min in blocking buffer (PBS with 0.2% Triton X-100 and 0.5% BSA). Incubation with primary antibody diluted in blocking buffer was performed overnight at 4°C. On the following day, ovaries were washed three times for 10 min in blocking buffer and incubated for 2-4 h at RT with secondary antibody diluted in blocking buffer. After washing the ovaries 3 times for 10 min with blocking buffer, ovaries were rinsed with PBS and mounted in DAPI Fluoromount-G (Thermo Fisher Scientific, OB010020). Samples were imaged with a Leica TCS SP8 with an HC PL APO CS2 63x/1.4 oil objective and Leica Application Suite X (LasX, Version 3.5.2.18963). Images were processed with FIJI v2.1.0/1.53c ^86^ and figures prepared in Inkscape 1.0.2 (e86c8708, 2021-01-15).

### The following antibodies were used

DSHB: mouse anti-Fas3 (7G10, 1:100), mouse anti-aop (8B12H9, 1:100), mouse anti-LamC (LC28.26, 1:100), mouse anti-eya (10H6, 1:50).

rabbit anti-GFP (Cell Signaling #2956, 1:1000), rabbit anti-dpERK (Cell Signaling #4370, 1:100), rat anti-RFP (ChromoTek 5F8, 1:1000), rat anti-dpn (abcam ab195173, 1:100), rabbit anti-bin (gift from M. Frasch, 1:200), rabbit anti-zfh1 (gift from J. Skeath, 1:500), rabbit anti-cas (gift from W. Odenwald, 1:1000), mouse anti-GFP 3E6 (Invitrogen #A-11120, 1:100).

Secondary antibodies were purchased from Invitrogen and applied in a 1:1000 dilution: goat anti-mouse 488 (A-11029), goat anti-mouse 568 (A-11031), goat anti-rabbit 488 (A-11034), goat anti-rabbit 568 (A-11036), goat anti-rat 488 (A-11073), goat anti-rat 568 (A-11077).

### Single-cell RNA-sequencing

Each genotype was sequenced in duplicates with a matched wildtype control. Single-cell solutions were prepared as previously reported ^12^. During dissection we enriched for non-vitellogenic stages using micro-scissors where phenotypes allowed for it. Sequencing was performed using the Chromium Single-Cell 3’ Reagent Version 2 Kit (10× Genomics) for dataset 1 and the Chromium Next GEM Single-Cell 3’LT Kit v3.1 (10× Genomics) for dataset 2. Sequencing was performed on Illumina HiSeq 2500 for dataset 1 and NovaSeq 6000 for dataset 2 according to the respective 10× Genomics manual.

### Bioinformatic analysis

For dataset 1 reads were aligned to the *Drosophila* reference genome (dmel_r6.19) using STAR v2.5.1b ^87^ and resulting bam files were processed with Cell Ranger. For dataset 2 we aligned to dmel_r6.19 using STARSolo ^88^. Post-processing was performed with Seurat 4.1.1 for dataset 1 and 5.1.0 for dataset 2 using R ^89^. We filtered out low quality cells based on the number of detected genes and percent of mitochondrial reads. For dataset 1 batch correction was performed with the Seurat wrapper ‘rliger’ and clustering performed in Seurat with a resolution of 0.1. Clusters were further sub-clustered were necessary to identify previously established cell populations. Multimodal reference mapping and cell cycle scoring were performed using Seurat. Note that reference mapping of dataset 2 to the dataset 1 control required a recalculation of the UMAP coordinates which resulted in slightly different UMAP coordinates. The set of genes used for cell cycle analysis can be found in Table S12. Regulons of the *w^1118^* datasets from dataset 1 were determined using SCENIC 1.3.1 and the cisTarget v8 motif collection mc8nr ^49^. Regulon activity was then determined across genotypes using the PercentageFeatureSet() command of Seurat. Differential abundance was tested using the R package implementation of Milo ^48^. Scales in Seurat expression plots and maps display the expression in log((UMI + 1/total UMI)×10^4^). All scripts are available upon request.

### Statistics and reproducibility

Statistical analyses were performed with R 2022.07.2 or Prism v7.0c. Graphs were plotted in R. Boxes in box plots show the median and interquartile range; lines show the range of values within 1.5× of the interquartile range. Error bars show the S.E.M. All images and quantifications are representative of at least three independent experiments. Single-cell datasets were performed in duplicates with matched wildtype controls. For dataset 1 a total of four and for dataset 2 a total of 2 wildtype controls were performed.

## Data availability

Single-cell RNA-sequencing datasets will be available at GEO after peer-reviewed publication. Raw data associated with quantifications is available from the Source Data file. The processed single-cell RNA-sequencing datasets are available upon request.

## Acknowledgements

We are thankful to M. Frasch, J. Skeath and W. Odenwald for sharing antibodies and D. Montell, T. Schüpbach, R. DasGupta and T. Harris for providing fly lines. We also thank the Bloomington Stock Center, the Vienna Drosophila Resource Center and the Developmental Studies Hybridoma Bank for many stocks and reagents used in this paper. We thank Meike Schneider for initial analysis of cell type specific signaling pathway activation and are grateful to Florian Finkernagel for the alignment of dataset 2. We also thank Sven Bogdan and Bogdan lab members for fruitful scientific discussions and comments on the manuscript.

## Author contributions

S.A. conducted fly genetic work, dissections and immunostaining, microscopical analysis and quantifications. A.S. provided support in dissections and immunostainings. J.M.P. performed initial phenotypic characterization and preparation of samples for single-cell RNA-sequencing of dataset 1. T.G.N. financed dataset 1. K.R. conceived the study and performed experimental work, bioinformatic analysis and preparation of samples for single-cell RNA-sequencing of dataset 2. K.R. and T.G.N. wrote the manuscript with input from all authors.

## Competing interests

The authors declare no competing interests.

## Supplementary Tables

**Table S1: Sequenced cells in dataset 1**

Table containing numbers of cells per genotype, dataset and cluster of dataset 1.

**Table S2: Expression profile of novel wildtype clusters**

Expression profile of wildtype young and mature stalk cells compared to other other wildtype clusters. Values were calculated using FindMarkers() on the *w^1118^*dataset.

**Table S3: Expression profile of novel clusters**

Expression profile of posterior polar cells, DNA damage cells, JNK high cells and Hh stalk-like clusters. Values were calculated using FindMarkers() on the entire dataset.

**Table S4: Expression profile of Hh^OE^ clusters**

Expression profile of Hh^OE^ clusters in comparison to the respective *w^1118^* cluster. Values were calculated using FindMarkers().

**Table S5: Expression profile of λtop^4.4^ clusters**

Expression profile of λtop^4.4^ clusters in comparison to the respective *w^1118^* cluster. Values were calculated using FindMarkers().

**Table S6: Expression profile of RasV12 clusters**

Expression profile of Ras^V12^ clusters in comparison to the respective *w^1118^* cluster. Values were calculated using FindMarkers().

**Table S7: Expression profile of hepCA clusters**

Expression profile of hep^CA^ clusters in comparison to the respective *w^1118^* cluster. Values were calculated using FindMarkers().

**Table S8: Expression profile of yki3SA clusters**

Expression profile of yki^3SA^ clusters in comparison to the respective *w^1118^* cluster. Values were calculated using FindMarkers().

**Table S9: Expression profile of armS10 clusters**

Expression profile of arm^S10^ clusters in comparison to the respective *w^1118^* cluster. Values were calculated using FindMarkers().

**Table S10: Wildtype regulons**

Regulons calculated by SCENIC from the *w^1118^* datasets of dataset 1.

**Table S11: Regulon expression**

Expression of high confidence regulon genes calculated from the *w^1118^* dataset per cell.

**Table S12: Cell cycle genes**

Table of genes used to score the cell cycle phase.

**Table S13: Sequenced cells in dataset 2**

Table containing numbers of cells per genotype, dataset and cluster of dataset 2.

**Table S14: Expression profile of Hh^OE^,Ras^V12^ clusters**

Expression profile of FSC, pFC and MB 2-3 Hh^OE^,Ras^V12^ clusters in comparison to the respective *w^1118^* cluster of dataset 2. Values were calculated using FindMarkers().

**Table S15: Expression profile of Hh^OE^,λtop^4.4^ clusters**

Expression profile of FSC, pFC and MB 2-3 Hh^OE^,λtop^4.4^ clusters in comparison to the respective *w^1118^* cluster of dataset 2. Values were calculated using FindMarkers().

---

**Supplementary Figure 1:**
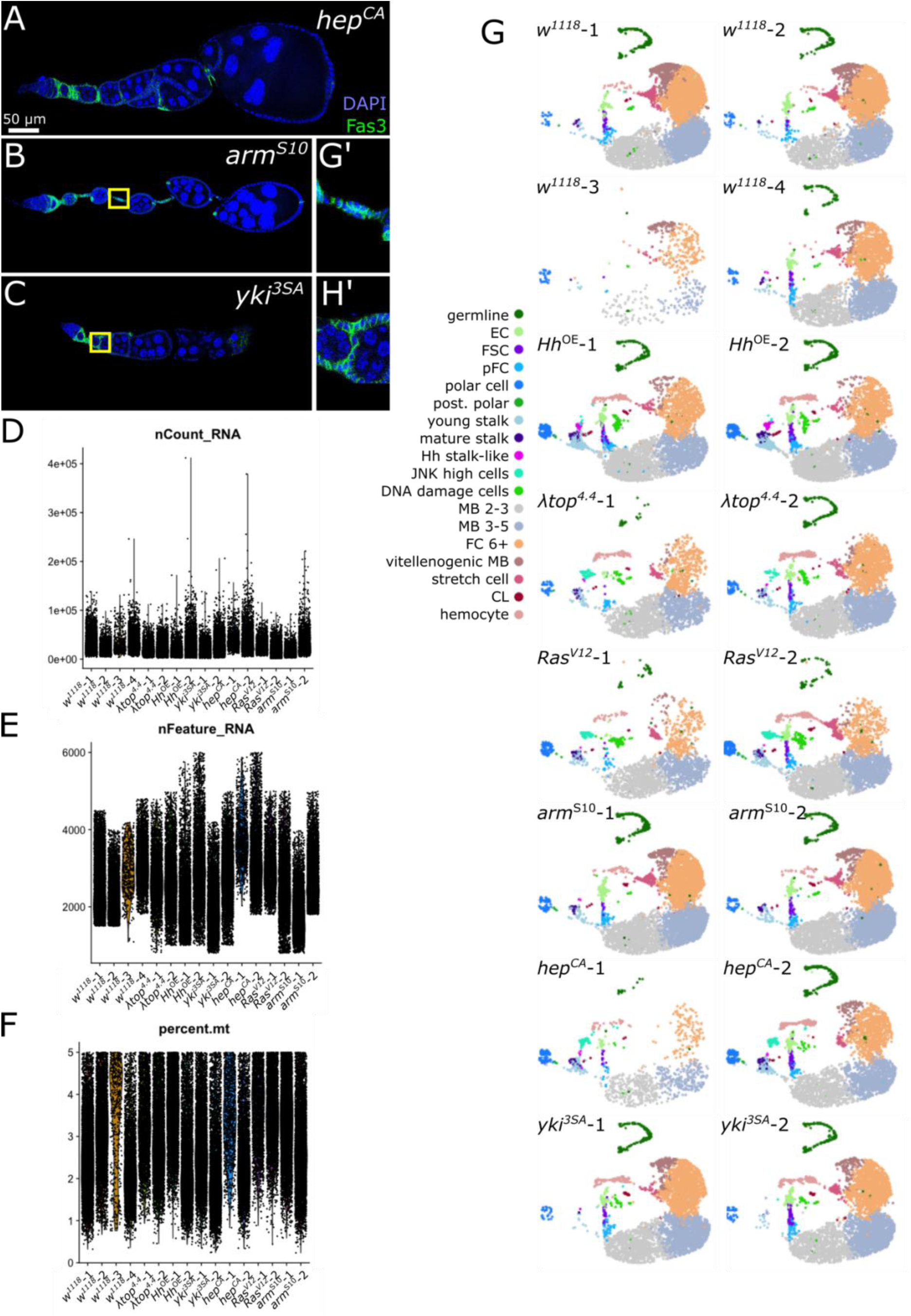
Quality controls and markers for cluster annotation. A-C) Ovarioles of 109-30^ts^ driving the respective genotypes. Insets in B) and C) show stalk regions. D-F) Quality features by dataset and per cell. D) Number of reads. E) Number of detected genes. F) Percent of mitochondrial reads. G) UMAP plot and cluster annotation of each dataset.

**Supplementary Figure 2:**
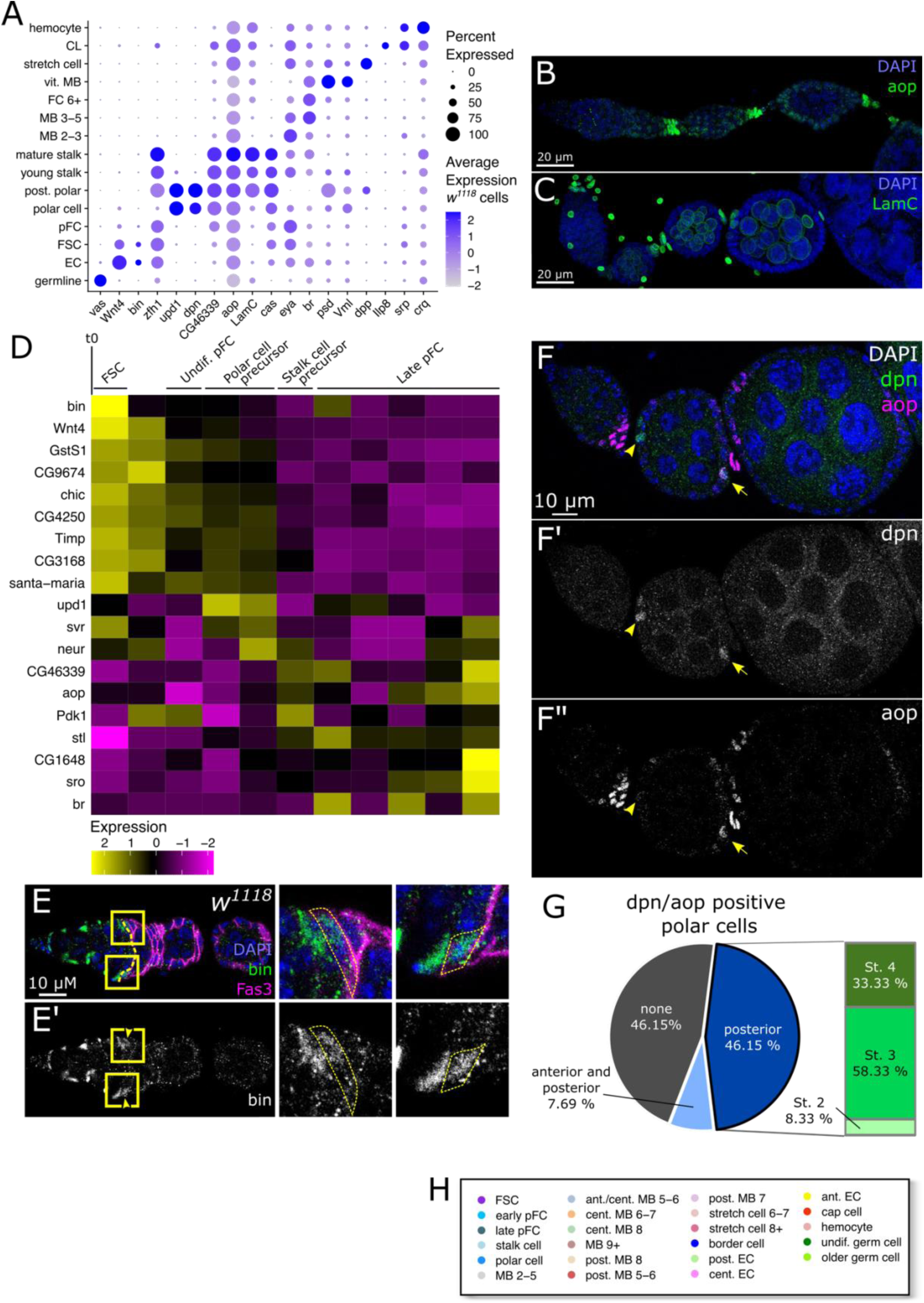
Bin and dpn are novel markers for FSCs and polar cells. A) Average and percent expression of cell type specific markers in *w^1118^*cells. Clusters that predominantly contained non-wildtype cells were excluded. B-C) Expression of the stalk cell markers in wildtype ovaries with the genotype *109-30^ts^* > *w^1118^* stained for DAPI (blue) and B) aop or C) LamC (green). D) Pseudotime analysis of wildtype FSCs and pFCs as reported in ^12^. *bin* marks FSCs at the beginning of pseudotime. E) Wildtype ovary of the genotype 109-30^ts^ > *w^1118^* stained for DAPI (blue), bin (green) and Fas3 (magenta). Note that a small number of Fas3 positive cells express bin (arrows). F) Expression of the polar cell marker dpn (green, white in F’) and the stalk cell marker aop (magenta, white in F”) in wildtype ovaries (*109-30^ts^* > *w^1118^*). DAPI (blue) marks nuclei. Young stages contain cells co-expressing dpn and aop (arrowheads). G) Quantification of dpn^+^/aop^+^ cells in n = 26 ovarioles. All double positive cells occupied positions corresponding to polar cells. H) Legend corresponding to Figure 2D).

**Supplementary Figure 3:**
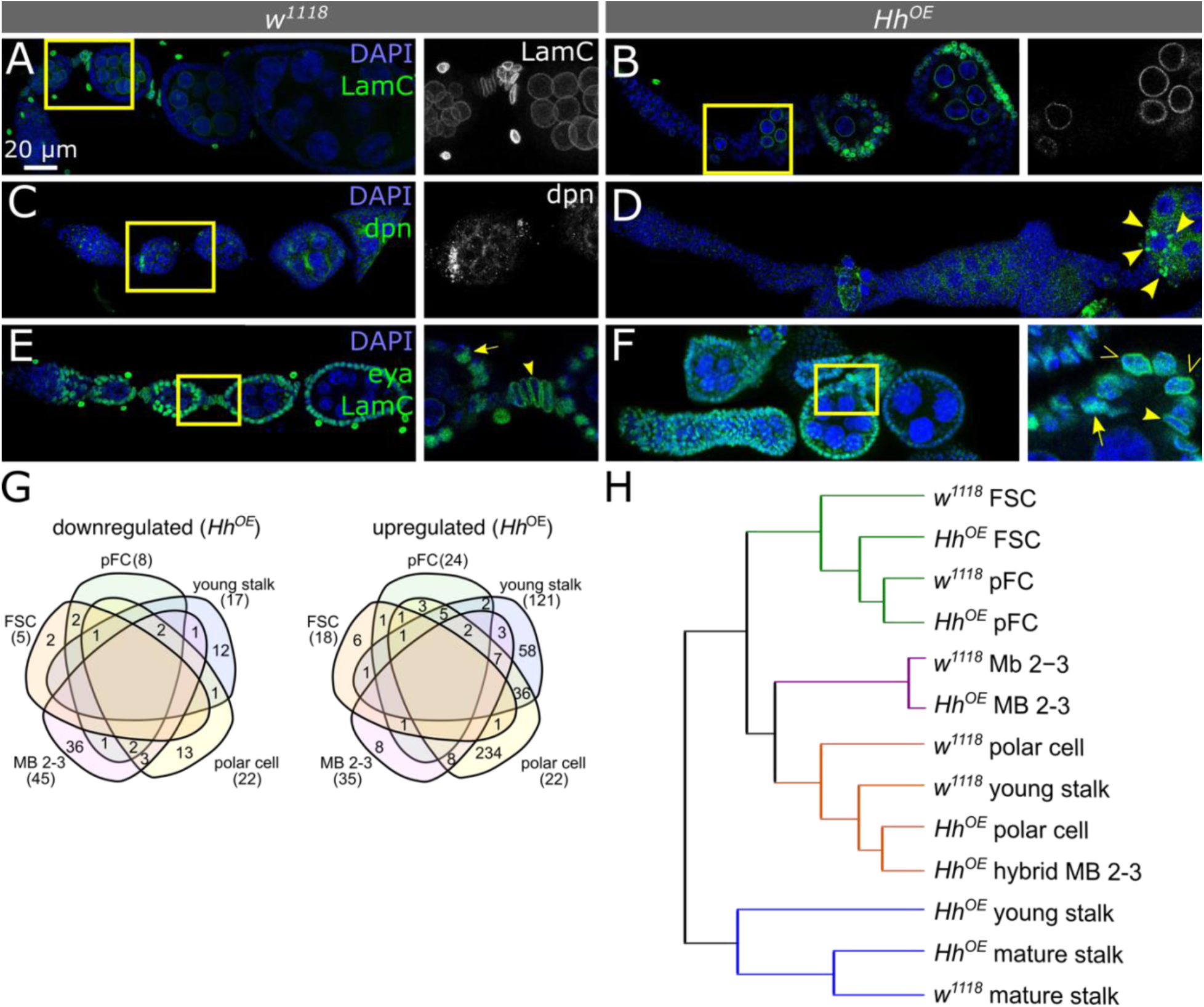
Hh overactivation affects follicle cell differentiation. A-F) Ovarioles of *109-30^ts^* crossed to *w^1118^*(A,C,E) or UAS-Hh^OE^ (B,D,F) stained for DAPI (blue) and cell type specific markers (green). A-B) LamC is expressed in control stalk cells once they matured. In early Hh^OE^ expressing follicle cells LamC is not expressed and becomes broadly expressed in later stages. C-D) dpn marks control polar cells and is not expressed in early stages of Hh^OE^ ovarioles. E-F) The MB marker eya and the stalk cell marker LamC are mutually exclusive in control ovarioles. Late stages of Hh^OE^ ovarioles display cell co-expressing LamC and eya (empty arrowhead) as well as cells positive for either LamC (arrowhead) or eya (arrow). G) Venn diagrams of down- and upregulated Hh target genes when compared to the *w^1118^* control of indicated clusters. Differentially expressed genes with a padj < 0.05 and log2FC ≥ 2 were considered. H) Cluster tree showing the relationship between *w^1118^*and Hh^OE^ clusters with 109-30-Gal4 activity. Hh^OE^ young stalk groups with mature stalk clusters, reminiscent of the early induction of mature stalk cell markers. Hybrid clusters (Hh^OE^ polar cells and Hh^OE^ MB 2-3 hybrid) group with *w^1118^* young stalk cells, implicating the induction of stalk markers in these clusters.The FSC and pFC clusters and MB 2-3 clusters which do not contain hybrid cells display relationships as expected.

**Supplementary Figure 4:**
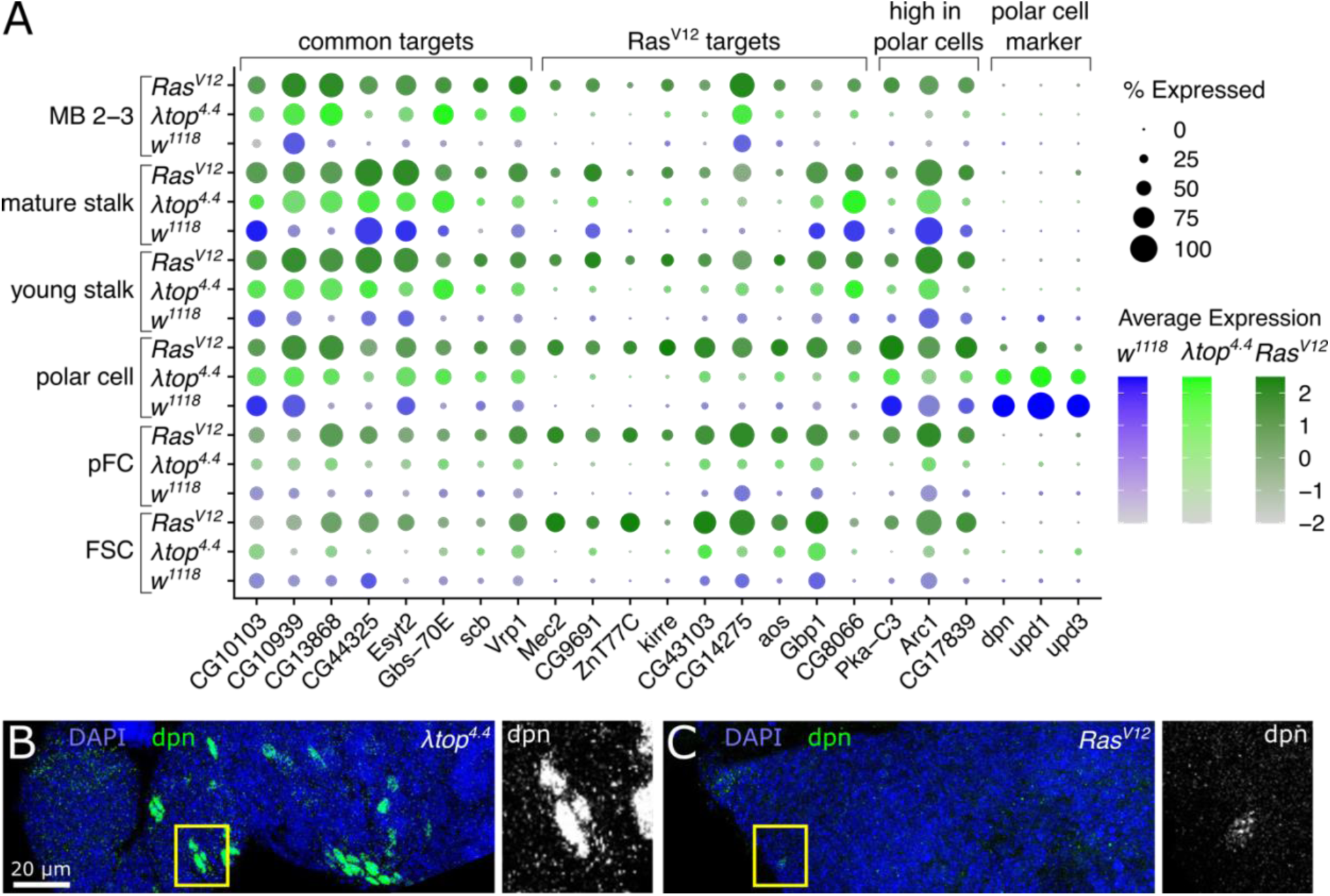
Ras^V12^ induces a distinct set of target genes. A) Average and percent expression of Ras^V12^ target genes. B-C) Ovarioles of *109-30^ts^*inducing λtop^4.4^ (B) or Ras^V12^ (C) stained for the nuclear marker DAPI (blue) and the polar cell marker dpn (green). Note that while induction of λtop^4.4^ results in clusters of dpn positive cells, Ras^V12^ results in a loss of dpn staining.

**Supplementary Figure 5:**
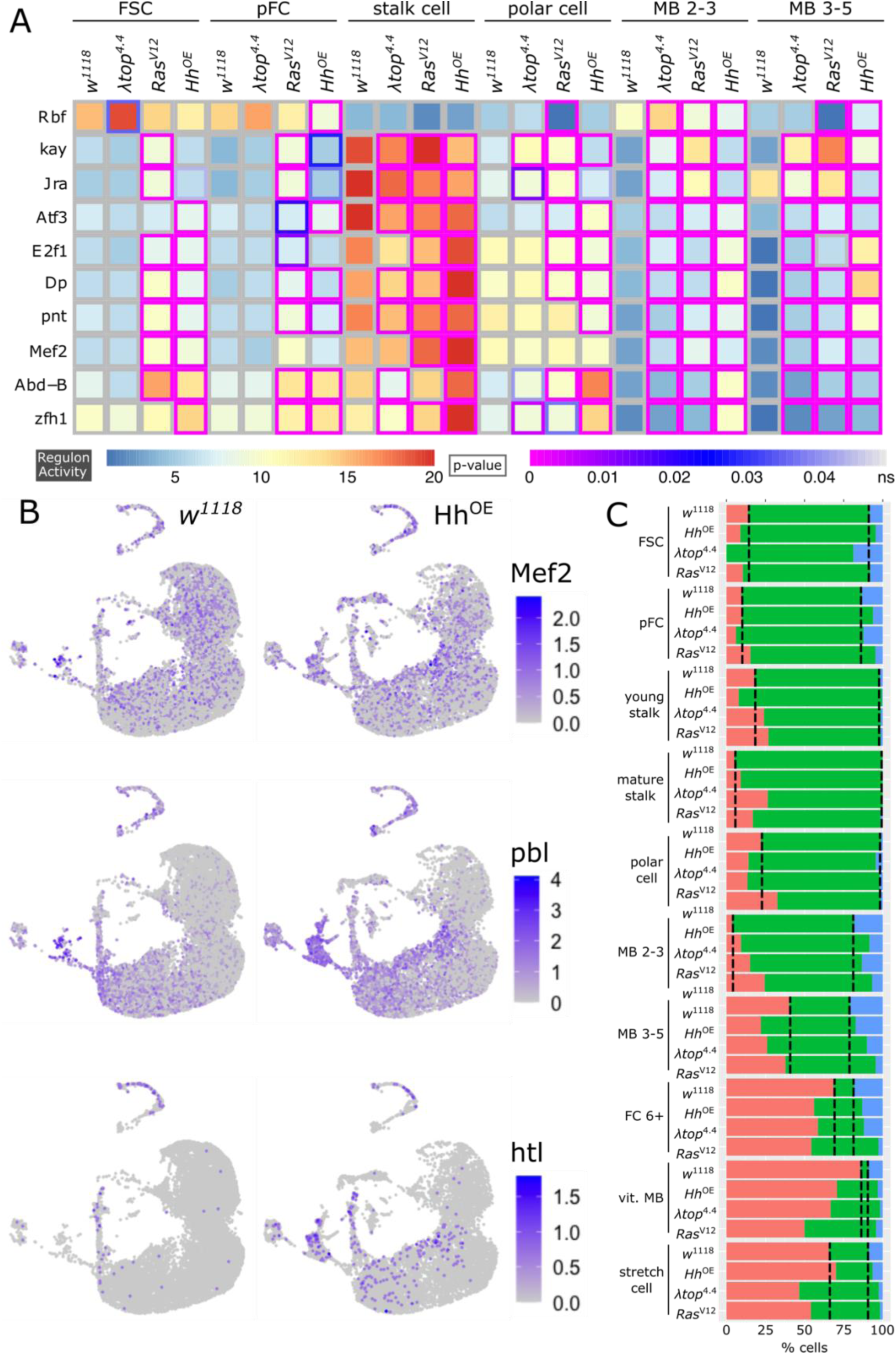
Signaling pathway perturbation affects transcription factor activity. A) Heatmap of regulon activity. Regulons with high confidence target genes were identified in the *w^1118^*dataset and expression was scored in datasets of all genotypes. Frame color marks significance of regulon activity change. B) UMAP plots of the wildtype *w^1118^* and the Hh^OE^ datasets for RNA expression of *Mef2*, *pbl* and *htl*. C) Bar graph of the percentages of cells per cluster in G1, G2M and S phase of the indicated genotypes.

**Supplementary Figure 6:**
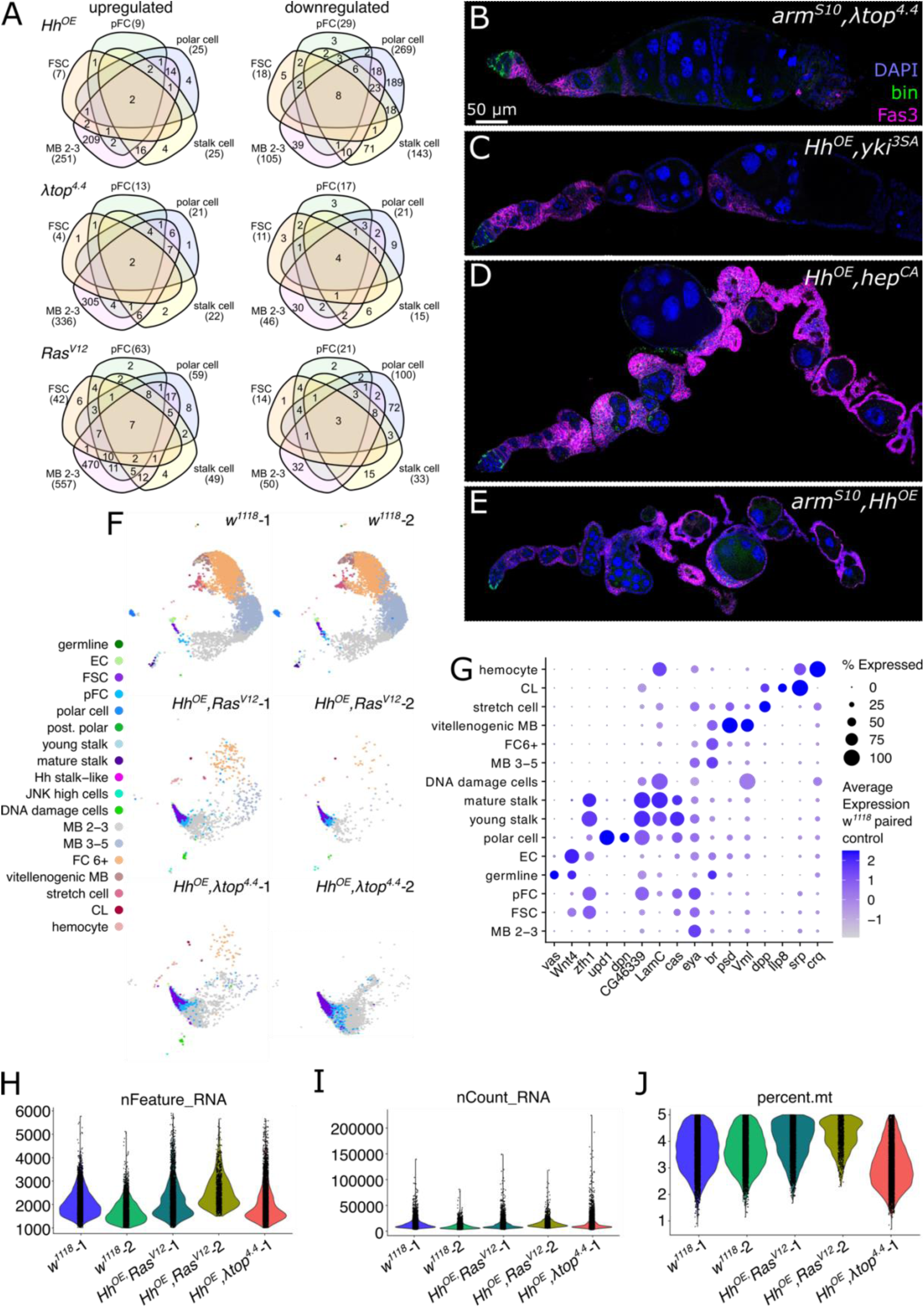
Tumorigenic follicle cell growth is specifically induced by overactivation of Hh^OE^ and λtop^4.4^ or Ras^V12^. A) Venn diagrams of differentially expressed genes with a padj value < 0.05 and a log2FC ≥ 2 per indicated genotype and cluster in comparison to the respective *w^1118^* cluster. B-E) Ovarioles of *109-30^ts^* driving double overactivation of B) arm^S10^,λtop^4.4^, C) Hh^OE^,yki^3SA^, D) Hh^OE^,hep^CA^ or E) arm^S10^,Hh^OE^ stained for DAPI (blue), Fas3 (magenta) and bin (green). F) UMAP plot showing reference mapping result of indicated genotypes. These *w^1118^* datasets were performed in paired experiments to the double overactivation datasets (dataset 2) and are different from the *w^1118^* datasets shown in Figure 1–5 (dataset 1). The Hh^OE^,λtop^4.4^-2 dataset was excluded from further analysis due to low data quality. G) Average and percent expression of cell type specific markers used for cluster annotation in cells of the paired *w^1118^* control. H-J) Violin plots showing the quality measures of detected features (H), number of counts (I) and percent of mitochondrial reads (J) per cell and dataset.

**Supplementary Figure 7:**
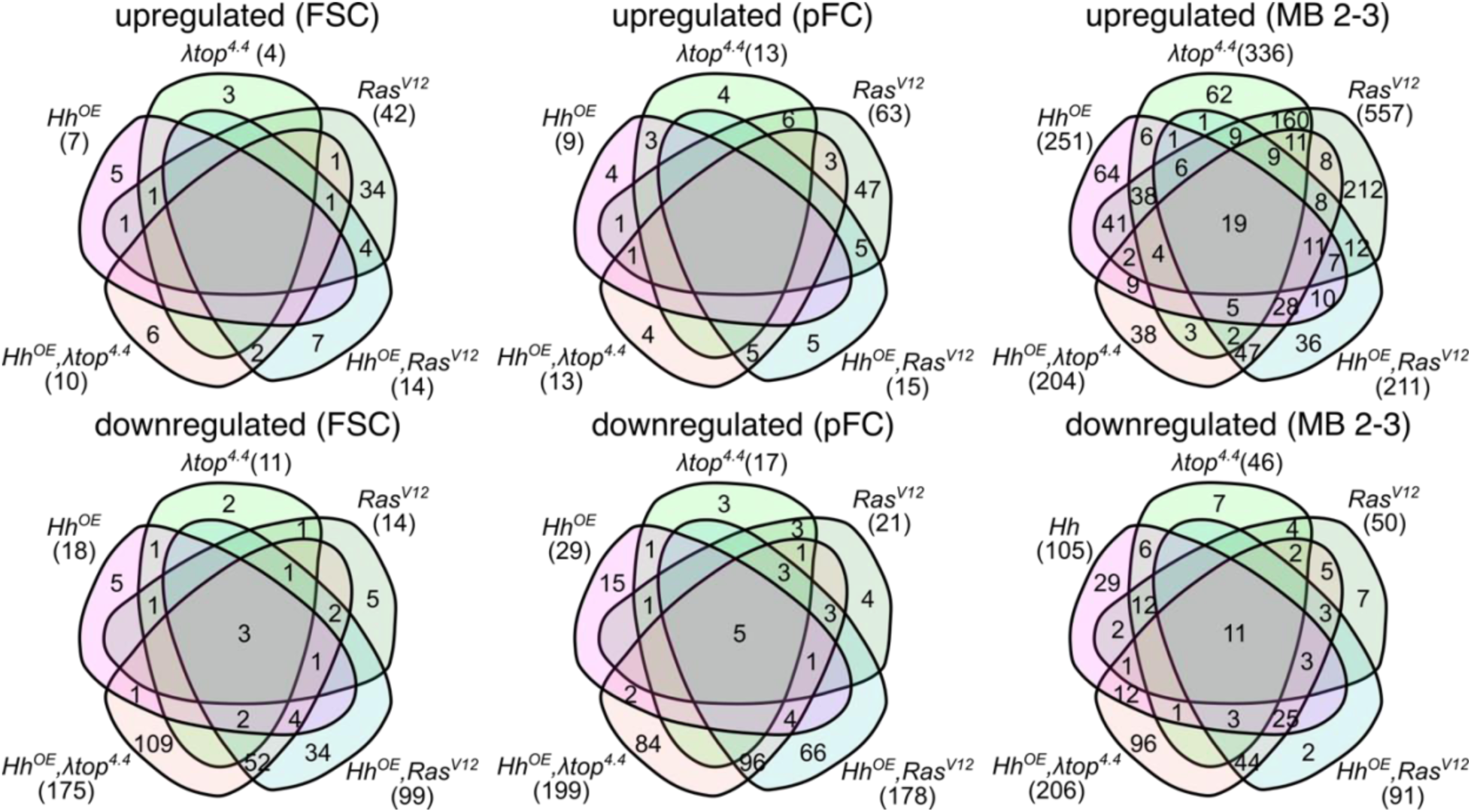
A majority of Hh-EGFR/Ras target genes are not misregulated by Hh or EGFR/Ras alone. Venn diagrams showing up- and downregulated genes with a p_adj_ value < 0.05 and a log2FC ≥ 2 in FSCs, pFCs and MB 2-3 clusters in different genotypes as indicated in comparison to the respective paired *w^1118^* control. Note that a large number of genes is specifically targeted only by double overactivation of Hh^OE^ and either λtop^4.4^ or Ras^V12^.

## Notes

### Competing Interest Statement

The authors have declared no competing interest.

### Summary of Updates

Added a relevant author, changed author order (agreed to by all authors)

### Funding

- NIH Common Fund: GM136348
- Röntgen-Behring Stiftung

